# Sensory Deprivation Independently Regulates Neocortical Feedforward and Feedback Excitation-Inhibition Ratio

**DOI:** 10.1101/283853

**Authors:** Nathaniel J. Miska, Leonidas M.A. Richter, Brian A. Cary, Julijana Gjorgjieva, Gina G. Turrigiano

## Abstract

Brief (2-3d) monocular deprivation (MD) during the critical period induces a profound loss of responsiveness within layer 4 of primary visual cortex (V1). This has largely been ascribed to long-term depression (LTD) at thalamocortical synapses onto pyramidal neurons, while a contribution from intracortical inhibition has been controversial. Here we used optogenetics to probe feedforward thalamocortical and feedback intracortical excitation-inhibition (E-I) ratios following brief MD. While thalamocortical inputs onto pyramidal neurons were depressed, there was stronger depression onto PV+ interneurons, which shifted the thalamocortical-evoked E-I ratio toward excitation. In contrast, feedback intracortical E-I ratio was shifted toward inhibition, and a computational model of layer 4 demonstrated that these opposing shifts produced an overall suppression of layer 4 excitability. Thus, feedforward and feedback E-I ratios onto the same postsynaptic target can be independently regulated by visual experience, and enhanced feedback inhibition is the primary driving force behind loss of visual responsiveness.

## INTRODUCTION

The fine-tuning of microcircuit function in primary sensory cortex requires sensory experience during an early critical period (Espinosa and Stryker, 2012; Gainey and Feldman, 2017; Hensch, 2005; Hubel and Wiesel, 1970), but the plasticity mechanisms that drive this refinement are not fully defined. In the visual system of rodents, the critical period extends from roughly postnatal days 21 - 35 (Espinosa and Stryker, 2012), when visual deprivation has catastrophic effects on visual function, including loss of visual responsiveness to the deprived eye (Frenkel and Bear, 2004), reduced visual acuity (Fagiolini et al., 1994), loss of tuning to many stimulus characteristics (Espinosa and Stryker, 2012), and a suppression of firing within primary visual cortex (V1) (Hengen et al., 2013; Hengen et al., 2016). Even very brief (2-3 days) monocular deprivation (MD) during the critical period induces a rapid loss of visual responsiveness to the deprived eye in both binocular (V1b) and monocular (V1m) regions of V1 (Frenkel and Bear, 2004, Kaneko et al., 2008). This rapid loss of deprived-eye responsiveness is thought to be primarily cortical in origin (Espinosa and Stryker, 2012; Gainey and Feldman, 2017; Sommeijer et al., 2017), but it remains controversial which forms of intracortical plasticity drive this loss of function. In particular, is it unclear whether classic Hebbian long-term depression (LTD) is the sole mediator (Heynen et al., 2003; Smith et al., 2009), or whether more complex changes in cortical circuitry that modify the relative recruitment of excitation and inhibition are equally or perhaps more important (House et al., 2011; Kuhlman et al., 2013; Li et al., 2014). Here we address this question by using optogenetic, electrophysiological, and modeling approaches to quantify the impact of brief MD on excitation-inhibition balance within defined feedforward and feedback pathways in V1m.

Traditionally, the loss of visual responsiveness after brief MD has been ascribed to LTD at excitatory synapses onto layer 4 or layer 2/3 pyramidal neurons (Crozier et al., 2007; Espinosa and Stryker, 2012; Frenkel and Bear, 2004; Heynen et al., 2003; Lambo and Turrigiano, 2013; Smith et al., 2009), including LTD at thalamocortical synapses onto excitatory neurons in layer 4 (Crozier et al., 2007; Khibnik et al., 2010). In this view, loss of responsiveness in layer 4 is a simple reflection of reduced efficacy at thalamocortical synapses onto excitatory postsynaptic neurons. In contrast, a number of recent studies have demonstrated that cortical inhibitory circuitry is also plastic and can be modified by brief MD (Kannan et al., 2016; Kuhlman et al., 2013; Maffei et al., 2006; Nahmani and Turrigiano, 2014; Sun et al., 2016). Given that the relationship between excitation (E) and inhibition (I) is critically important for neocortical information processing (Isaacson and Scanziani 2011, Haider and McCormick 2009), E-I ratio is likely a more relevant measure of circuit excitability than excitation alone (House et al., 2011; Li et al., 2014; Xue et al., 2014). However, it is currently unknown whether brief MD alters E-I ratio within specific microcircuit motifs in V1.

Primary sensory neocortical regions such as V1 have a stereotyped microcircuit architecture (Douglas et al., 1989; Van Hooser, 2007). Sensory information is relayed to cortex via thalamocortical inputs, which excite both pyramidal neurons and inhibitory interneurons within layer 4, forming a ‘feedforward’ circuit. In addition, pyramidal neurons recurrently excite other pyramidal neurons and interneurons within layer 4, forming an intracortical ‘feedback’ circuit. While these two circuits are intimately related and share both excitatory and inhibitory elements, they ultimately provide separable contributions to sensory-evoked cortical activity (Reinhold et al., 2015). Furthermore, ‘feedback’ recurrent inputs greatly outnumber feedforward thalamocortical inputs (Ahmed et al., 1994; Bopp et al., 2017; Douglas et al., 1989), suggesting that the layer 4 feedback circuit is poised to disproportionately affect network excitability. The extent to which the feedforward thalamocortical and feedback intracortical components of synaptic drive onto the same postsynaptic neuron may be independently tuned by experience-dependent plasticity remains an unexplored question.

To characterize changes in thalamocortical versus intracortical circuits following brief MD, we used optogenetic methods to isolate and independently probe these respective circuit elements in layer 4 of V1m. First, we expressed channelrhodopsin-2 (ChR2) in thalamocortical afferents in layer 4 and recorded from layer 4 pyramidal neurons after brief MD. Surprisingly, although thalamocortical inputs were indeed depressed by brief MD as expected (Khibnik et al., 2010), the E-I ratio was shifted toward excitation rather than inhibition; this effect was mediated by a disproportionate depression of thalamocortical synaptic strength onto layer 4 PV+ interneurons compared to neighboring pyramidal neurons. In contrast, probing intracortical E-I ratio through sparse expression of ChR2 in layer 4 pyramidal neurons revealed that this local feedback circuit was significantly shifted toward inhibition following brief MD. We modeled these opposite shifts in feedforward versus feedback E-I ratio and found that changes in feedback E-I ratio outweighed changes in feedforward E-I ratio, producing an overall depression in network activation over a wide range of parameter space. These data suggest that contrary to the prevailing view, brief MD amplifies rather than suppresses feedforward thalamocortical inputs onto layer 4 pyramidal neurons, and the loss of responsiveness within layer 4 arises instead through reduced intracortical amplification by the local feedback circuit. Further, our data establish that feedback and feedforward E-I ratio onto the same postsynaptic target can be independently adjusted by visual experience.

## RESULTS

In rodents, brief MD during the critical period decreases firing rates (Hengen et al., 2013, 2016) and reduces visual responsiveness in deprived V1 (Frenkel and Bear, 2004; Kaneko et al., 2008; Khibnik et al., 2010). The extent to which this reflects thalamocortical LTD (Crozier et al., 2007; Smith et al, 2009) versus reorganization of E-I networks (Kuhlman et al., 2013; Maffei et al., 2006) remains controversial. To examine this, we used optogenetics to probe and compare the relative strengths of excitation and inhibition in thalamocortical feedforward and intracortical feedback circuits within layer 4. For all experiments, control and deprived data were obtained from the same animals, with deprived data obtained from V1m contralateral to the deprived eye, and control data obtained from V1m ipsilateral to the deprived eye.

### Brief MD induces LTD-like depression at thalamocortical synapses onto pyramidal neurons in V1m

Brief visual deprivation reduces the ability of thalamocortical synapses to evoke visual responses in V1 (Frenkel and Bear, 2004; Khibnik et al., 2010) and occludes the induction of LTD at synapses onto principle neurons in layer 4 (Crozier et al., 2007). To determine whether brief MD induces an absolute reduction in thalamocortical postsynaptic strength, we developed a protocol that allowed us to probe quantal amplitudes specifically at thalamocortical synapses. We selectively labeled primary thalamocortical afferents in V1 with a ChR2-mCherry fusion protein by stereotaxically injecting adeno-associated virus (AAV-ChR2-mCherry) bilaterally into the dorsal lateral geniculate nucleus (dLGN) of the thalamus of Long-Evans rat pups (Figure 1A, Kloc and Maffei, 2014; Wang et al., 2013). Following a 1.5 – 2 wk period to allow expression and transport of ChR2, we observed dense mCherry-positive thalamocortical axon terminals in V1, with strongest expression in layer 4 as expected (Figure 1B,C). Next, we developed an optogenetically-evoked, desynchronized vesicle release paradigm to measure quantal amplitudes at labeled thalamocortical synapses. We obtained whole-cell recordings from layer 4 pyramidal neurons in the presence of tetrodotoxin (TTX) to block action potentials, 4-aminopyradine (4-AP) to enhance the excitability of presynaptic terminals (Petreanu et al., 2009), and (2*R*)-amino-5-phosphonovaleric acid (AP5) and picrotoxin (PTX) to isolate AMPA-mediated mEPSCs. Under these conditions, long (1 - 5 s) pulses of blue laser light elevated mEPSC frequency to (on average) 485 ± 48% of baseline (Figure 1E, Figure S1A, P < 0.0001). At moderate stimulus intensities, this allowed us to measure evoked quantal events from thalamocortical synapses with minimal contamination from spontaneous mEPSCs.

**Figure 1:**
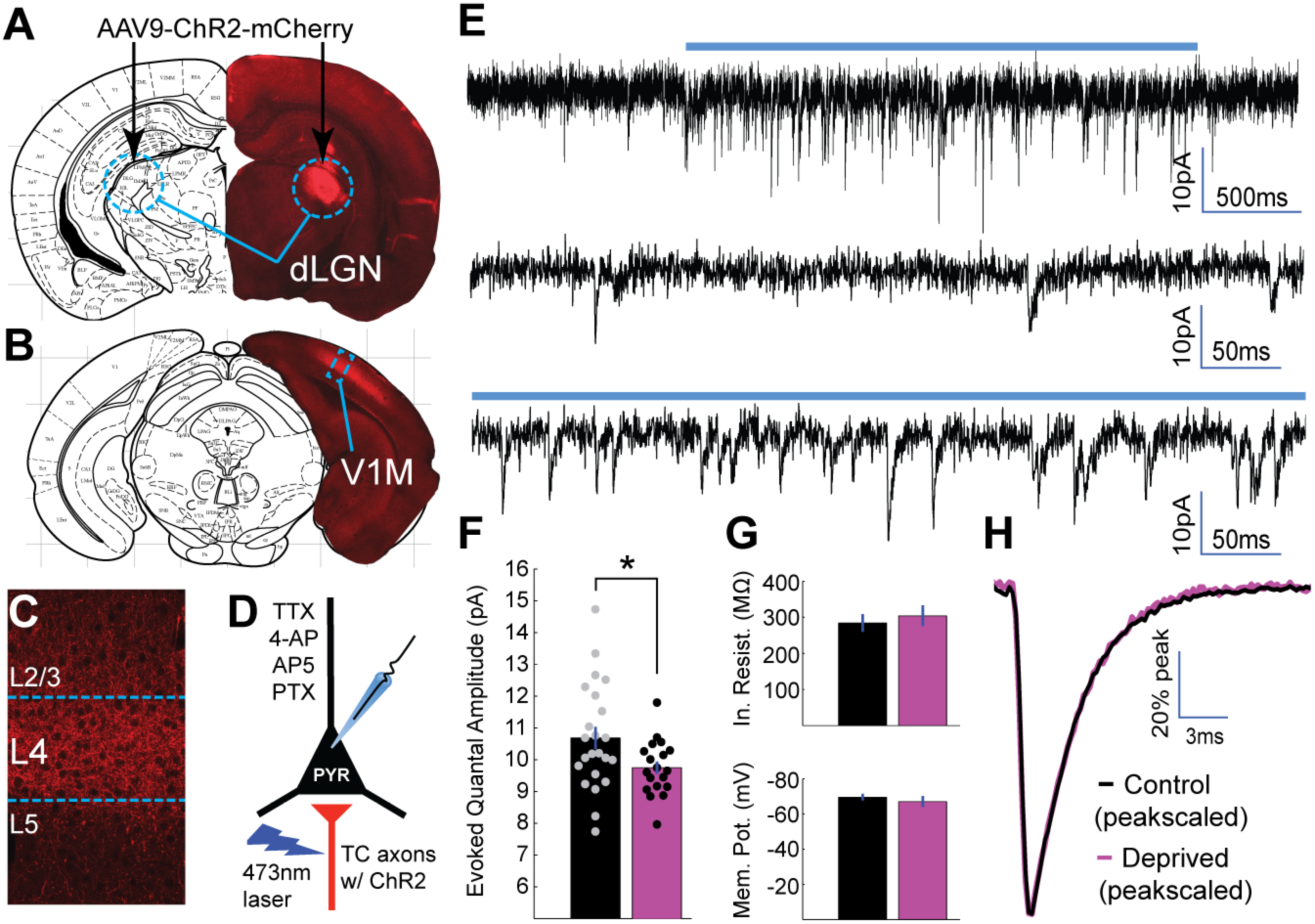
Brief MD reduces thalamocortical quantal amplitude onto layer 4 pyramidal neurons. **A**) Stereotaxic injection of AAV-ChR2-mCherry into dLGN (red) **B**) leads to reporter expression within thalamocortical axons in V1, **C**) with densest innervation in layer 4 (L4). **D**) Whole-cell recordings were obtained from layer 4 pyramidal neurons in a drug cocktail of TTX, 4-AP, AP5, and PTX to isolate excitatory quantal events while stimulating local ChR2-mCherry+ thalamocortical (TC) axons (red) with 473 nm light. **E**) Top: example recording of quantal events during 2 s laser stimulation (blue bar). Middle: Pre-stimulus spontaneous mEPSCs. Bottom: Evoked quantal events during laser stimulation. **F**) Mean evoked quantal amplitudes for control (black) and deprived (magenta) neurons (control n = 22, deprived n = 18, P = 0.016, 2-sample t-test). **G**) Mean input resistance (top) and resting membrane potential (bottom) for control and deprived neurons. **H**) Overlaid peak-scaled evoked event waveform averages for control (black) and deprived (magenta) neurons, to illustrate kinetics. For all bar plots here and below, circles represent individual values, and error bars indicate ± SEM.

Examination of the amplitude distributions of evoked events revealed a similar distribution to spontaneous events, with no multimodal peaks indicative of contamination by multiquantal events (Neubig et al., 2003; Figure S1B). The mean amplitude and kinetics of evoked thalamocortical quantal events were similar to spontaneous quantal events, as expected (Gil et al., 1999, Figure S1C). Brief MD induced a modest but significant reduction in evoked thalamocortical quantal amplitude (Figure 1F, control = 10.7 pA ± 0.35 pA, deprived = 9.8 pA ± 0.20 pA, P < 0.05), without any impact on input resistance, resting membrane potential (Figure 1G), waveform kinetics (Figure 1H), or spontaneous mEPSC amplitudes (Figure S1D). This reduction in quantal amplitude following brief MD is consistent with the induction of a postsynaptic depression at thalamocortical synapses onto layer 4 pyramidal neurons.

If this reduction in thalamocortical quantal amplitude reflects the induction of LTD, then further LTD induction should be occluded. To test this, we adapted a low frequency LTD induction paradigm (Crozier et al., 2007) for optogenetic stimulation, which involved activating thalamocortical axons repeatedly at 1 Hz for 10 min paired with brief postsynaptic depolarization to −50 mV in voltage clamp. This paradigm reliably induced sustained depression of evoked thalamocortical EPSCs onto control neurons (Figure 2A,B,E,G, 63.1 ± 7.2% of baseline amplitude). In contrast, deprived neurons exhibited significantly less depression following LTD induction (Figure 2B,F,G, 89.3 ± 9.7% of baseline amplitude, P < 0.05). There were no changes in passive properties following LTD induction in either condition (Figure 2C,D). Taken together with the change in thalamocortical quantal amplitude, this strongly suggests that brief MD induces LTD-like depression of thalamocortical synapses onto layer 4 pyramidal neurons in deprived V1m.

**Figure 2:**
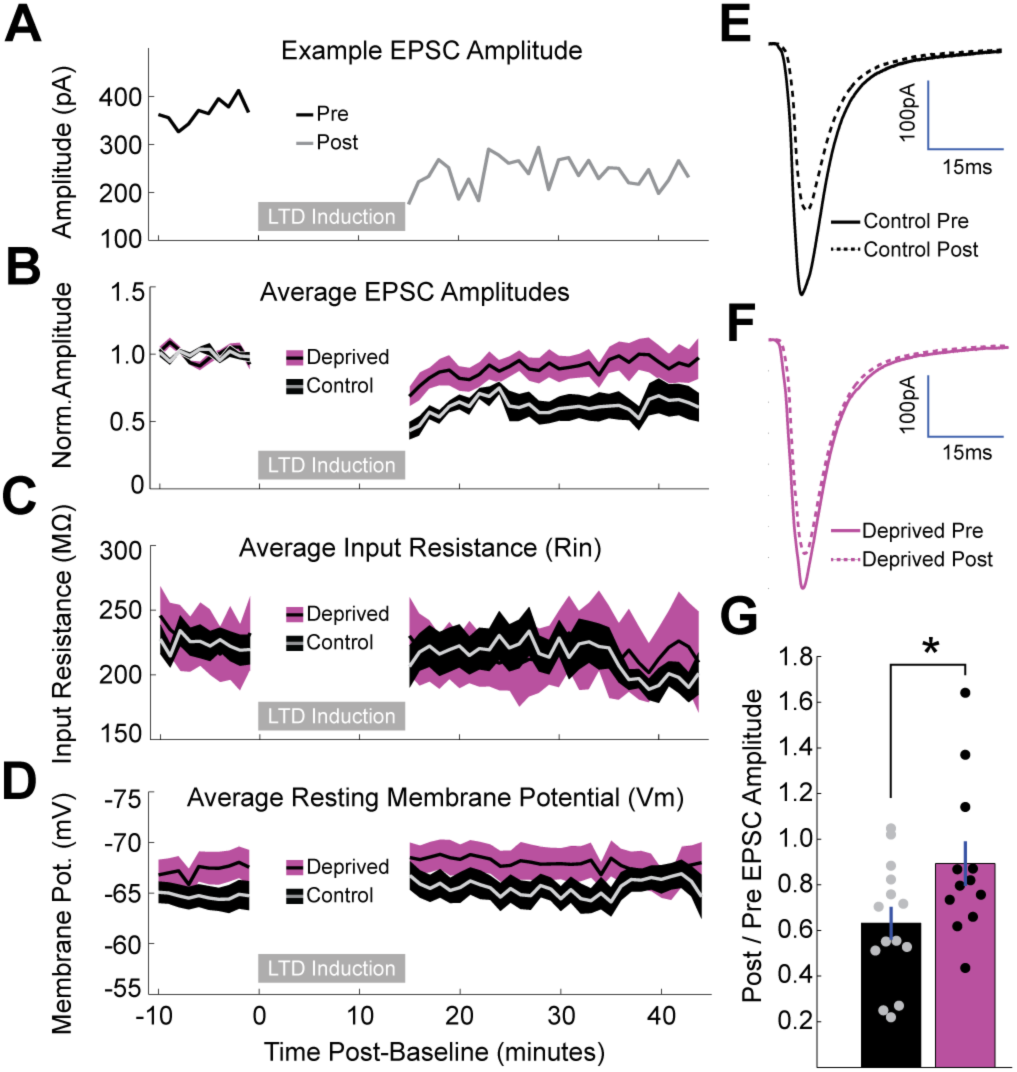
Brief MD occludes induction of LTD at thalamocortical synapses onto layer 4 pyramidal neurons. **A**) Representative example of LTD induction from a nondeprived neuron, with pre-induction evoked EPSC amplitudes plotted in black, post-induction amplitudes plotted in grey, and LTD induction period represented as a grey bar. **B**) Mean, baseline-normalized evoked EPSCs plotted for control (grey with black shading) and deprived (black with magenta shading) neurons. Shaded region represents SEM. **C**) Mean input resistance (Rin) and **D**) mean resting membrane potential (Vm) plotted for control and deprived neurons. **E,F**) Baseline-normalized EPSC waveform averages for control neurons (**E**) pre-induction (black, solid) and post-induction (black, dashed), and deprived neurons (**F**) pre-induction (magenta, solid) and post-induction (magenta, dashed). **G**) Mean post/pre EPSC amplitude ratio for control (black) and deprived (magenta) neurons (control n = 14 cells, deprived n = 12 cells, P = 0.0374, 2-sample t-test).

### Thalamocortical-evoked E-I ratio shifts toward excitation following brief MD

Thalamocortical inputs to layer 4 target GABAergic interneurons as well as pyramidal neurons, and thalamocortical activation evokes a mix of monosynaptic excitation and disynaptic inhibition onto layer 4 pyramidal neurons (Cruikshank et al., 2010; Kloc and Maffei, 2014). If, as previously suggested (Crozier et al., 2007; Espinosa and Stryker, 2012; Khibnik et al., 2010; Smith et al., 2009), LTD at thalamocortical to pyramidal synapses is the major driver of the loss of visual responsiveness in layer 4, then the ratio of thalamocortical-evoked excitation to thalamocortical-evoked inhibition should shift to favor inhibition after brief MD. Layer 4 and layer 2/3 inhibitory synapses can also exhibit experience-dependent plasticity (House et al., 2011; Maffei et al., 2006; Nahmani and Turrigiano, 2014; Xue et al., 2014), but how visual experience impacts thalamocortical E-I ratio has not been assessed.

E-I ratio in layer 2/3 has been shown to exhibit significantly less variability than excitation or inhibition alone (Xue et al., 2014), suggesting it represents a conserved measure regulated at the level of the neuron or local microcircuit. We measured feedforward thalamocortical E-I ratio using brief (2 ms) pulses of blue laser light to activate thalamocortical afferents while holding postsynaptic layer 4 pyramidal neurons at the experimentally-determined reversal potential for excitation or inhibition (Figure 3A,B). Paired monosynaptic EPSCs versus disynaptic IPSCs scaled linearly at different stimulus intensities (Figure 3B, inset), indicating that thalamocortical E-I ratio is conserved as additional thalamocortical axons are recruited. Paradoxically, even though there was depression at thalamocortical to pyramidal neuron synapses following brief MD (Figures 1 and 2), the thalamocortical-evoked E-I ratio shifted significantly to favor excitation in deprived neurons in both C57BL/6J mice (Figure 3C, control E-I ratio = 0.094 ± 0.010, deprived E-I ratio = 0.148 ± 0.016, P < 0.01) as well as in Long-Evans rats (Figure S3). There was no change in passive properties between deprived and control cells (Figure 3D).

**Figure 3:**
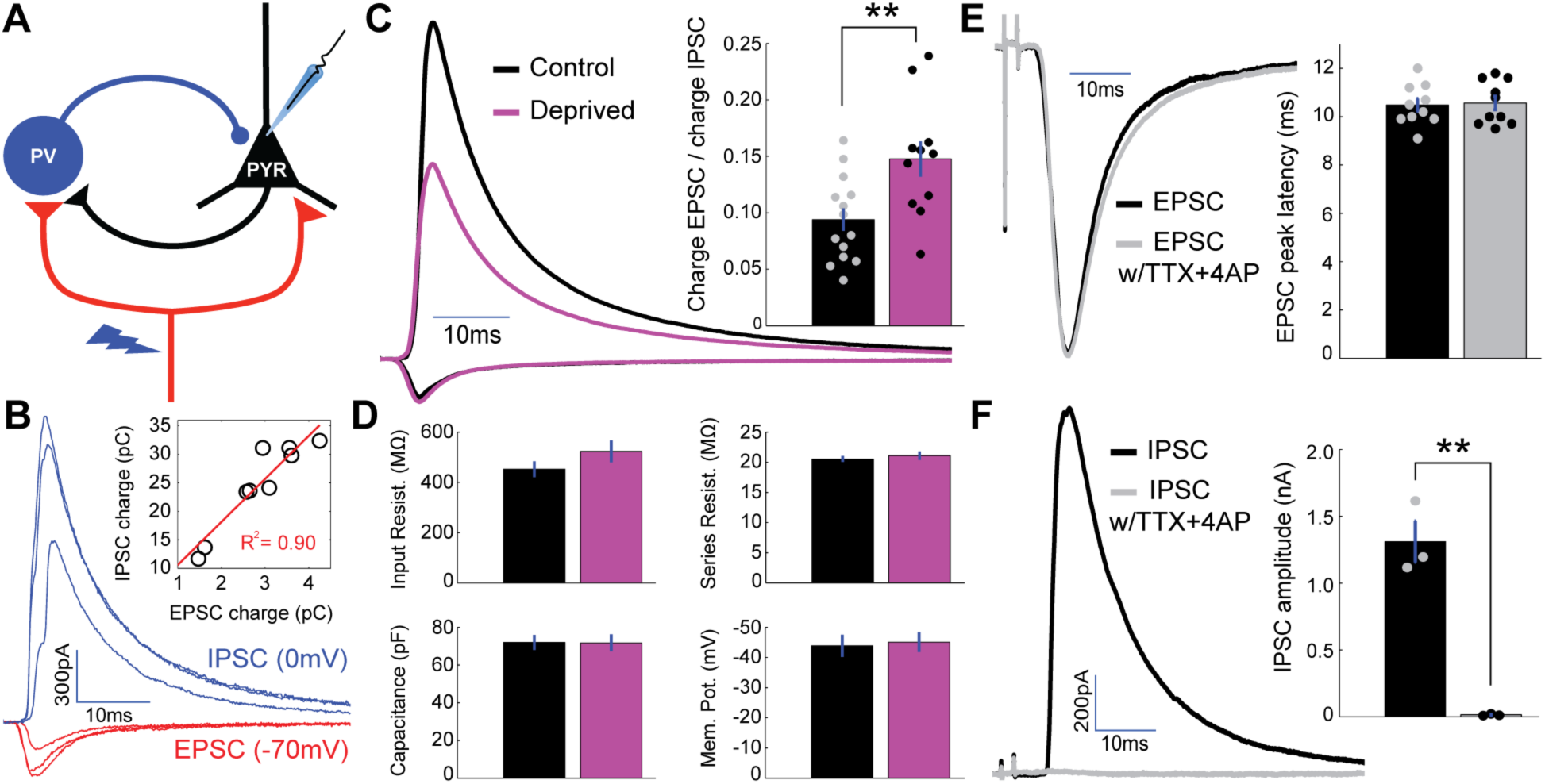
Brief MD increases thalamocortical-evoked Excitation-Inhibition (E-I) ratio. **A**) Layer 4 pyramidal neurons were voltage-clamped in normal ACSF while stimulating local ChR2-mCherry+ thalamocortical axons with 473 nm light. **B**) Representative example traces from a single neuron): brief (2 ms) stimuli were given at increasing laser intensity while alternating between the experimentally determined inhibitory and excitatory reversal potentials (−70 mV & 0 mV, respectively) in order to record paired monosynaptic thalamocortical-evoked EPSCs and disynaptic thalamocortical-evoked IPSCs. Inset: EPSC charge plotted versus IPSC charge for individual E-I pairs from this example neuron, with linear fit plotted in red (R^2^ = 0.90). **C**) Mean EPSC charge-normalized EPSC and IPSC traces for control (black) and deprived (magenta) neurons. Inset: Mean E-I charge ratio for control (black) and deprived (magenta) neurons (control n = 14, deprived n = 11, P = 0.0063, 2-sample t-test). **D**) Mean input resistance, series resistance, capacitance, and resting membrane potential for control (black) and deprived (magenta) neurons. **E**) Mean peak-scaled thalamocortical EPSCs evoked in regular ACSF (black) and following perfusion of TTX+4-AP (grey). Bar plot: EPSC peak latency in regular ACSF (black) and following perfusion of TTX+4-AP (grey) (n = 10, P = 0.82, 2-sample t-test). **F**) Mean thalamocortical IPSCs evoked in regular ACSF (black) and following perfusion of TTX+4-AP (grey). Bar plot: IPSC amplitude in regular ACSF (black) and following perfusion of TTX+4-AP (grey) (n = 3, P = 0.0014, 2-sample t-test).

To verify that we were not recruiting polysynaptic excitation with this stimulation paradigm, we used perfusion of TTX+4-AP to assess whether evoked thalamocortical EPSCs were monosynaptic (Cruikshank et al., 2010; Ji et al., 2016; Petreanu et al., 2009). At the stimulus intensities used, evoked thalamocortical EPSC kinetics and peak latency were unchanged following perfusion of TTX+4-AP (Figure 3E, control peak latency = 10.5 ± 0.27 ms, TTX+4-AP peak latency = 10.6 ± 0.28 ms, P = 0.82), while IPSCs were completely abolished (Figure 3F, control peak amplitude = 1320 ± 164pA, TTX+4-AP peak amplitude = 15 ± 4 pA, P < 0.01). Hence, evoked EPSCs represent monosynaptic thalamocortical inputs, and evoked IPSCs represent disynaptic inputs from local inhibitory interneurons driven by thalamocortical activation. Given that excitatory thalamocortical synaptic strength is depressed following brief MD (Figures 1 and 2), this implies that there is a concurrent and even greater weakening of thalamocortical-recruited feedforward inhibition onto pyramidal neurons. Notably, this change in thalamocortical E-I ratio is in the opposite direction from that needed to account for the well-documented reduction in visual responsiveness induced by brief MD.

### Enhanced depression at thalamocortical synapses onto PV+ interneurons following brief MD

One possibility that could explain a shift in thalamocortical-evoked E-I ratio toward excitation (Figure 3) is that brief MD induces depression at thalamocortical synapses onto *both* pyramidal neurons and GABAergic interneurons, but the relative decrease is *greater* onto GABAergic interneurons. The two major subtypes of GABAergic interneurons in layer 4 are parvalbumin (PV)+ and somatostatin (SST)+ interneurons (Ji et al., 2016; Pfeffer et al., 2013; Rudy et al., 2011). PV+ interneurons receive much stronger thalamic drive than SST+ interneurons (Beierlein et al., 2003; Cruikshank et al., 2010; Ji et al., 2016; Urban-Ciecko and Barth, 2016; Yavorska and Wehr, 2016), suggesting that they mediate the vast majority of thalamocortical-evoked feedforward inhibition. To target PV+ interneurons, we crossed mice carrying PV-Cre (Hippenmeyer et al., 2005) and Rosa26-STOP-tdTomato (Ai9, Madisen et al., 2010) alleles, such that progeny express tdTomato in PV+ interneurons. To verify that layer 4 PV+ interneurons in V1 fire in response to thalamocortical activation in the range of stimulus intensities used to assess E-I ratio (Figure 3B), we recorded in current clamp from tdTomato-expressing PV+ interneurons and nearby pyramidal neurons in layer 4 while stimulating thalamocortical axons. Indeed, PV+ interneurons fired readily even to modest thalamocortical stimulation and with a much lower threshold than pyramidal neurons (Figure S4, P < 0.0001), suggesting that these neurons contribute substantially to the feedforward E-I measurements.

To test whether thalamocortical synapses onto PV+ interneurons are depressed more than thalamocortical synapses onto pyramidal neurons, we measured relative thalamocortical drive to both cell types using paired recordings in tandem with thalamocortical stimulation (Figure 4A,B). Evoked thalamocortical EPSCs onto PV+ interneurons were generally larger with faster kinetics than EPSCs onto pyramidal neurons as expected (Figure 4C; Hull et al., 2009; Kloc and Maffei, 2014), and EPSC amplitude onto the two cell types scaled linearly with stimulus intensity (Figure 4D). After brief MD, evoked thalamocortical EPSCs onto PV+ interneurons were significantly smaller relative to paired neighboring pyramidal neurons (Figure 4E), resulting in an increase in thalamocortical-evoked pyramidal/PV EPSC charge ratio (Figure 4F, control ratio = 0.260 ± 0.027, deprived ratio = 0.369 ± 0.034, P < 0.05). These data show that brief MD induces a greater depression of thalamocortical synapses onto layer 4 PV+ interneurons than onto neighboring pyramidal neurons; this in turn should reduce the ability of thalamocortical stimulation to recruit PV-mediated inhibition onto layer 4 pyramidal neurons and thus is predicted to contribute to an increase in thalamocortical-evoked E-I ratio.

**Figure 4:**
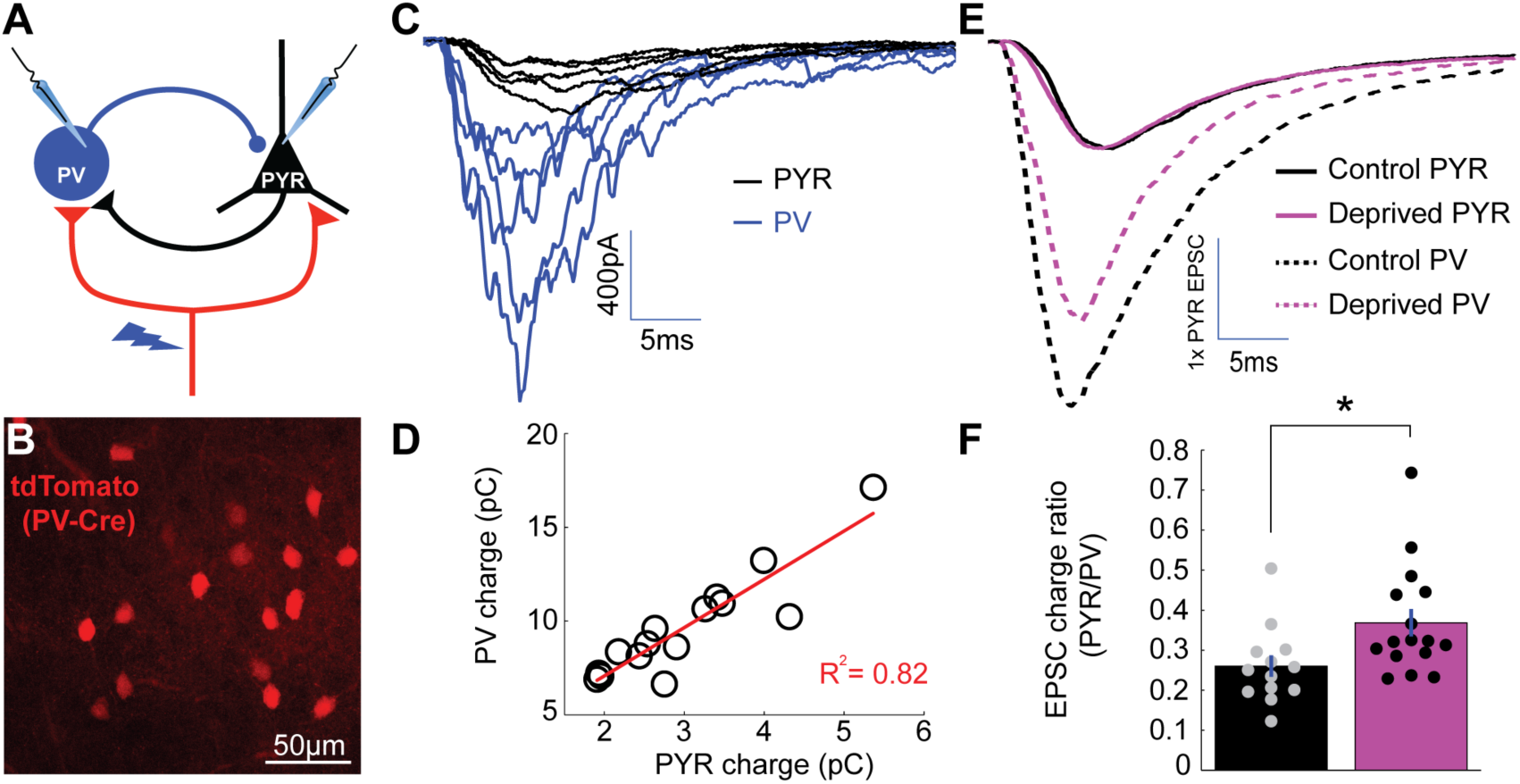
Enhanced depression at thalamocortical synapses onto PV+ interneurons following brief MD. **A**) Whole-cell recordings were obtained from pairs of pyramidal neurons (black) and nearby PV+ interneurons (blue) while stimulating local ChR2-mCherry+ thalamocortical axons (red). **B)** PV+ interneurons were targeted by reporter (tdTomato) expression. **C**) Representative example traces from a pyramidal and PV pair while stimulating thalamocortical axons at a range of stimulus intensities, with pyramidal traces in black (PYR) and PV+ interneuron traces in blue (PV). Note the larger thalamocortical EPSCs in the PV+ interneuron compared to the pyramidal neuron. **D**) Charge of thalamocortical EPSCs in the pyramidal neuron plotted against corresponding thalamocortical EPSCs in the PV+ interneuron. Linear fit plotted in red (R^2^ = 0.82). **E**) Averaged traces from all pairs (control in black and deprived in magenta) normalized to pyramidal EPSC peak amplitude. **F**) Evoked thalamocortical EPSC charge ratio (PYR/PV) for control (black) and deprived (magenta) pairs (control n = 13 pairs, deprived n = 16 pairs, P = 0.0103, Wilcoxon rank sum test).

### Brief MD does not affect intrinsic excitability of layer 4 neurons or inhibitory strength from PV+ interneurons to pyramidal neurons

In addition to changes at thalamocortical synapses, relative changes in intrinsic excitability of PV+ interneurons and pyramidal neurons could potentially contribute to the increase in thalamocortical E-I ratio induced by brief MD. We measured intrinsic excitability of layer 4 pyramidal neurons and PV+ interneurons, targeted using the reporter line as described above, by generating firing rate versus current (FI) curves under conditions where synaptic inputs were blocked (Figure 5). There was no significant difference in firing rate between deprived and control PV+ interneurons at any current injection (Figure 5B). Furthermore, there was no difference in input resistance or rheobase (Figure 5C). Similarly, pyramidal intrinsic excitability, input resistance, and rheobase were unaffected by brief MD (Figure 5D-F).

**Figure 5:**
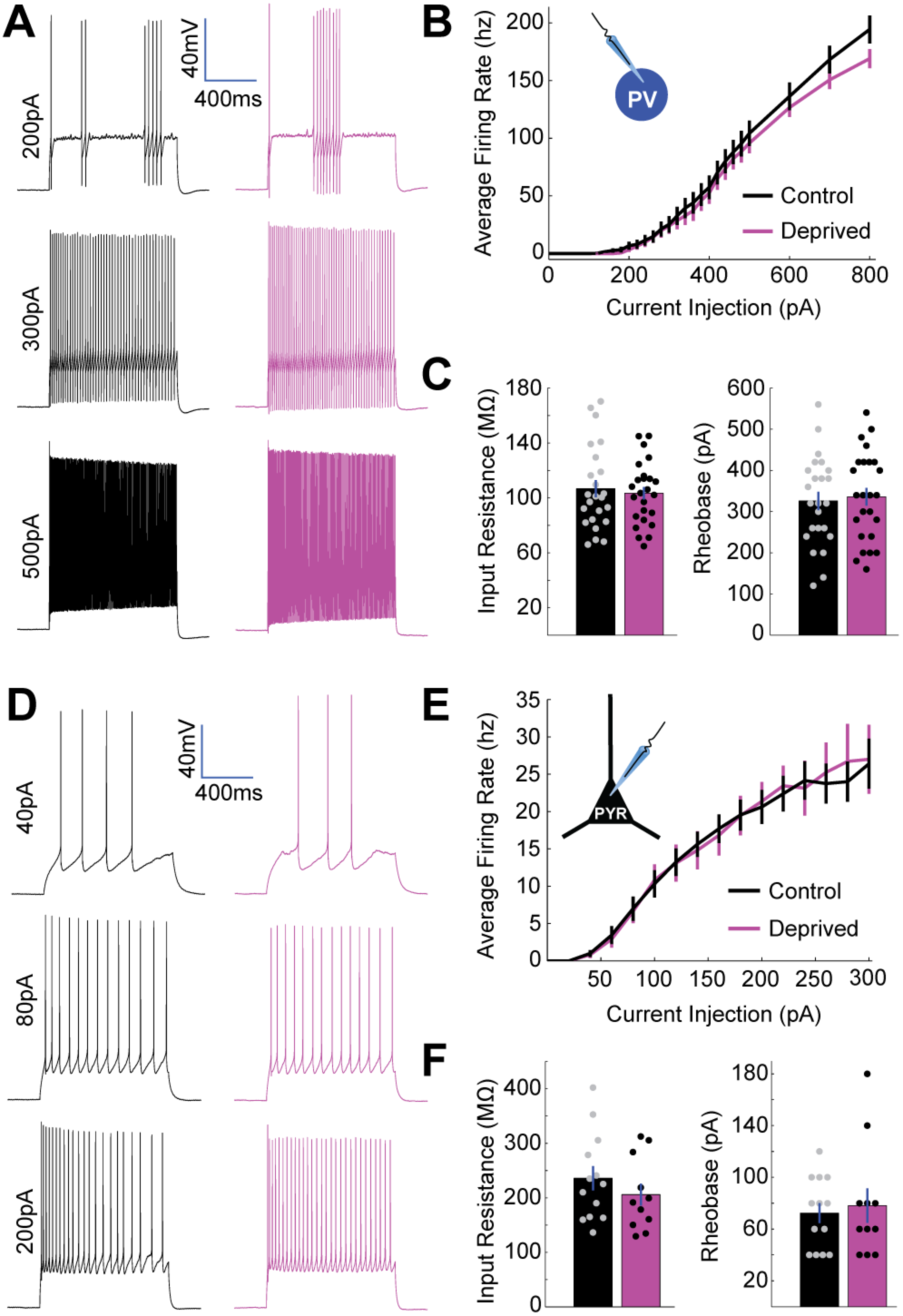
No change in intrinsic excitability of layer 4 PV+ interneurons or pyramidal neurons following brief MD. **A**) Representative example traces from a control PV+ interneuron (black) and a deprived PV+ interneuron (magenta) showing firing responses to 1 s long current injections of 200 pA (top), 300 pA (middle), and 500 pA (bottom). **B**) PV+ interneuron firing rate versus current injection (FI) plotted for all neurons in control (black, n = 24) or deprived (magenta, n = 25) conditions. **C**) Mean input resistance (left) and mean rheobase (right) for control (black) and deprived (magenta) PV+ interneurons. **D**) Representative example traces from control (black) and deprived (magenta) layer 4 pyramidal neurons showing firing responses to 1 s long current injections of 40 pA (top), 80 pA (middle), and 200 pA (bottom). **E**) Pyramidal neuron firing rate versus current injection (FI) plotted for all neurons in control (black, n = 13) or deprived (magenta, n = 11) conditions. **F**) Mean input resistance (left) and mean rheobase (right) for control (black) and deprived (magenta) pyramidal neurons.

Weakening of inhibitory synaptic strength from PV+ interneurons onto layer 4 pyramidal neurons is another mechanism that could contribute to the increased E-I ratio following brief MD. To test this possibility, we developed an optogenetic minimal stimulation paradigm to obtain putative unitary IPSCs from individually targeted PV+ interneurons onto nearby layer 4 pyramidal neurons. For this experiment, we crossed mice carrying PV-Cre and Rosa26-STOP-ChR2-EYFP (Ai32, Madisen et al., 2012) alleles, such that progeny express ChR2-EYFP in PV+ interneurons. While recording from layer 4 pyramidal neurons, nearby individual reporter-expressing PV+ interneurons were targeted with a low-intensity focal laser spot (Figure 6A), and laser intensity was iteratively increased until IPSCs were elicited by ∼50% of stimuli (Figure 6B). This procedure was repeated for up to 4 nearby PV+ interneurons while recording from up to 3 layer 4 pyramidal neurons at a time, allowing for a comparatively high throughput assessment of putative unitary IPSC amplitudes. Inhibitory connection strength varied markedly across connections, even from the same presynaptic PV+ interneuron (Figure 6B), with a small population of very strong connections (Figure 6D). By patching onto the same PV+ interneuron we had stimulated optogenetically, we could show that spike-evoked unitary IPSCS and optogenetically evoked putative unitary IPSCs were indistinguishable (Figure 6C), indicating that this approach generates reliable estimates of unitary connection strength.

**Figure 6:**
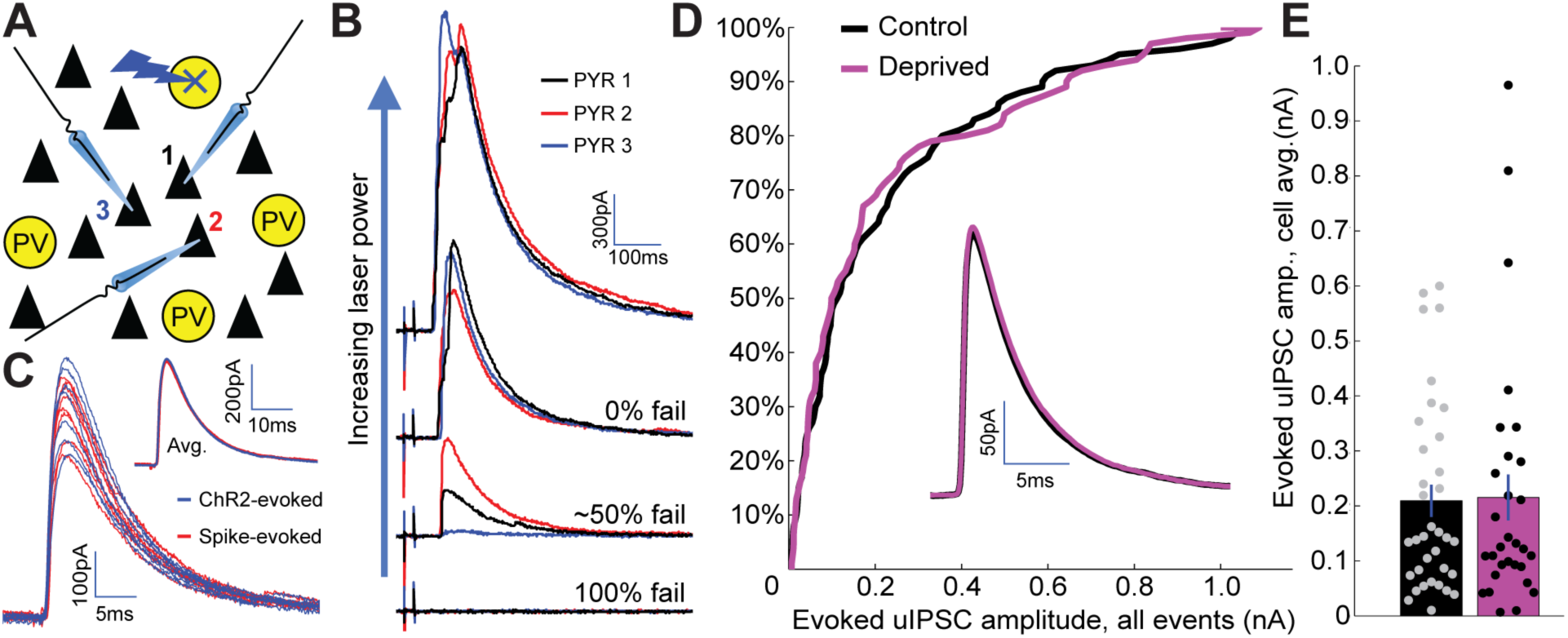
No change in PV+ interneuron to pyramidal neuron inhibitory synaptic strength following brief MD. **A**) Recording/stimulating configuration. While recording from multiple pyramidal neurons, nearby PV+ interneurons expressing ChR2-YFP were targeted serially with focal laser stimulation. **B**) For each targeted PV+ interneuron, laser stimulation strength was initially very low and was gradually increased until one or more recorded pyramidal neurons exhibited ∼50% rate of failure in evoked IPSCs. **C**) Example IPSCs evoked using optogenetic stimulation (blue) or by patching the same PV+ interneuron and evoking individual spikes with current injection (red). Inset: average traces from each method overlaid. **D**) Cumulative probability distribution for all putative unitary IPSC amplitudes (includes multiple inputs per pyramidal neuron) is not significantly different between control and deprived conditions (control n = 68, deprived n = 59, P = 0.90, two-sample Kolmogorov-Smirnov test). Inset: average IPSC waveforms (cell-averages) are indistinguishable between control and deprived pyramidal neurons. **E**) Quantification of cell-averaged putative unitary IPSC amplitudes shows no significant difference between control and deprived pyramidal neurons (control amplitude = 209 ± 29 pA n = 35 neurons, deprived amplitude = 215 ± 42 pA n = 30 neurons, P = 0.73, Wilcoxon rank sum test).

Comparing optogenetically-evoked putative unitary IPSC amplitude between control and deprived neurons revealed no significant difference in mean amplitude (control amplitude = 209 ± 29 pA, deprived amplitude = 215 ± 42 pA, P = 0.73), or in the distribution of IPSC amplitudes (P = 0.90), between conditions (Figure 6D,E). Hence, changes in unitary synaptic strength from PV+ interneurons to pyramidal neurons do not significantly contribute to the weakening of thalamocortical-recruited inhibition following brief MD. Taken together, our data suggest that the major factor that increases feedforward thalamocortical E-I ratio onto pyramidal neurons is the disproportionate reduction in thalamic drive to PV+ interneurons.

### Local intracortical-evoked E-I ratio shifts toward inhibition following brief MD

It is well established that 2-3 d of MD reduces visual responsiveness (Kaneko et al., 2008), signal propagation (Wang et al., 2011), and firing rates in V1m (Hengen et al., 2013, 2016). Loss of visual responsiveness following brief MD has largely been ascribed to depression at thalamocortical synapses (Espinosa and Stryker, 2012; Khibnik et al., 2010; Smith et al., 2009), yet here we show that the thalamocortical-evoked E-I ratio onto layer 4 pyramidal neurons is increased, rather than reduced, by brief MD. One explanation for how layer 4 excitability decreases despite this increase in thalamocortical E-I ratio is that E-I ratio is regulated differently in feedforward thalamocortical circuitry and recurrent feedback intracortical circuitry. Since intracortical excitatory synapses outnumber thalamocortical synapses by a factor of ∼5 in neocortical layer 4 (Bopp et al., 2017) and by at least a factor of 2 onto PV+ interneurons (Kameda et al., 2012) a change in intracortical E-I ratio could potentially outweigh changes in thalamocortical circuitry to suppress excitability.

To measure intracortical E-I ratio in layer 4, we modified our optogenetic paradigm to allow us to label and activate a subset of layer 4 pyramidal neurons (Figure 7A) while recording from nearby uninfected pyramidal neurons. To achieve this we injected a low volume of AAV-FLEX-ChR2-tdTomato into V1M of Scnn1a-Tg3-Cre mice (Madisen et al., 2010), resulting in a fraction of layer 4 pyramidal neurons expressing ChR2-tdTomato (Figure 7B,C). Intracortical-evoked EPSCs and IPSCs were recorded in voltage clamp at the experimentally verified reversal potentials for inhibition and excitation. As observed for thalamocortical-evoked E-I events, EPSCs versus IPSCs scaled linearly as a function of stimulus intensities (Figure 7D, inset). Following perfusion of DNQX, IPSCs were abolished (Figure 7E, baseline IPSC amplitude = 909 ± 57 pA, DNQX IPSC amplitude = 12.2 ± 3.8 pA, P < 0.0001), indicating that ChR2-tdTomato expression was restricted to excitatory neurons. After brief MD, intracortical-evoked E-I ratios were significantly shifted toward greater inhibition in deprived versus control pyramidal neurons (Figure 7F, control ratio = 0.091 ± 0.009, deprived ratio = 0.066 ± 0.005, P < 0.05), with no accompanying change in passive properties (Figure 7G). Interestingly, intracortical and thalamocortical E-I ratios were indistinguishable under control conditions, but diverged dramatically after brief MD (Figure 7H, P < 0.0001, thalamocortical E-I values re-plotted from Figure 3C for comparison). This suggests that baseline E-I ratios are strongly conserved across feedforward and feedback microcircuits in layer 4 but can be independently regulated by brief visual deprivation.

**Figure 7:**
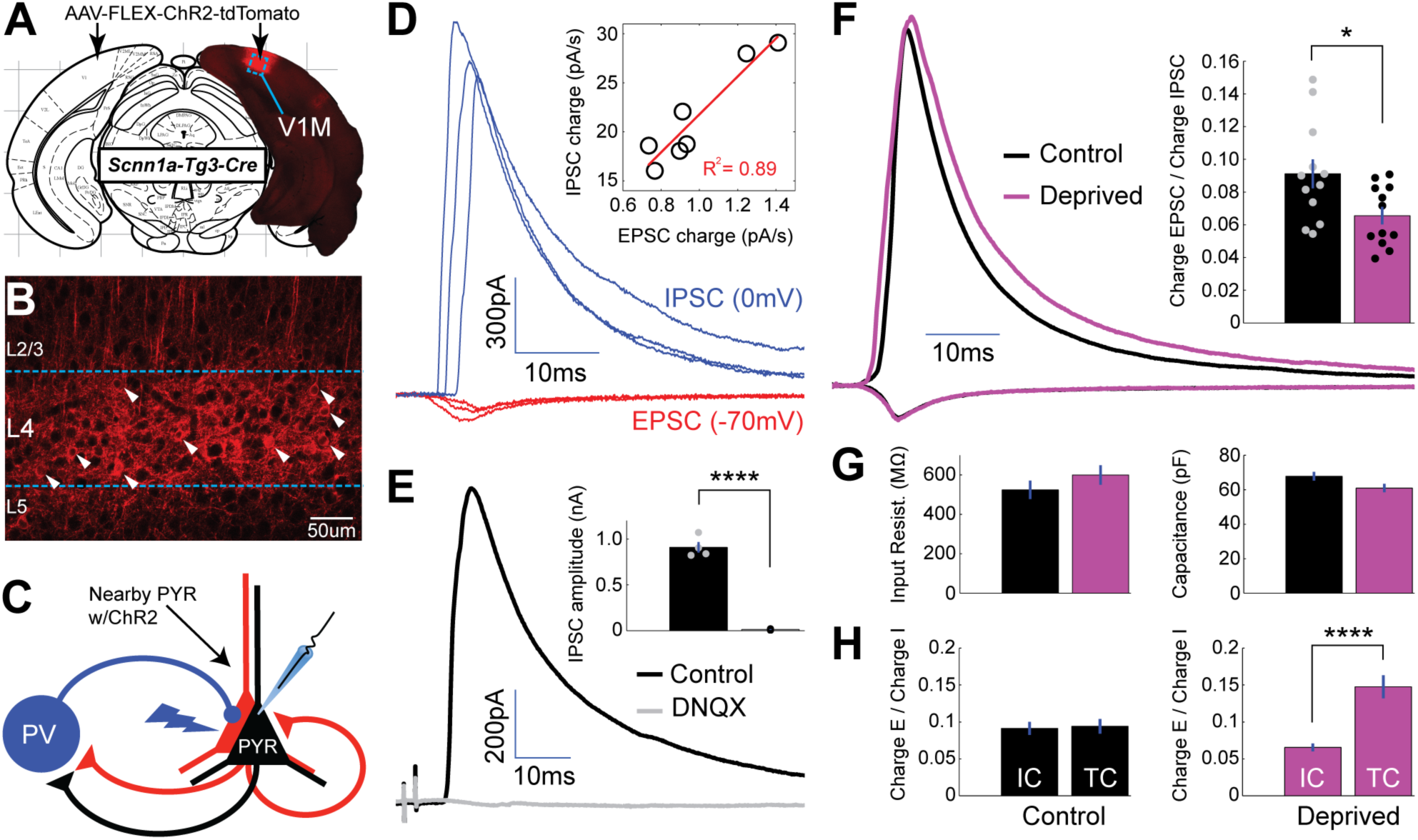
Local intracortical-evoked E-I ratio shifts toward inhibition following brief MD. **A**) Cre-dependent ChR2-tdTomato expression (red) from stereotaxic injection of AAV into V1M of Scnn1a-Tg3-Cre mice. **B**) Confocal image from site of viral infection, with infected layer 4 pyramidal neurons marked with white arrows. **C**) Diagram of recording configuration: whole-cell recordings were obtained from uninfected layer 4 pyramidal neurons while stimulating neighboring ChR2-tdTomato+ pyramidal neurons with brief (2 ms) pulses of 473 nm light. **D**) Representative example paired EPSCs (red) and IPSCs (blue) recorded at experimentally verified reversal potentials (−70 mV and 0 mV) at several stimulus intensities. Inset: EPSC versus IPSC charge elicited by a range of stimulus intensities for a single neuron, with linear fit plotted in red (R^2^ = 0.89). **E**) Evoked IPSCs in regular ACSF (black) and following perfusion of DNQX (grey) (n = 4). **F**) Mean EPSC charge-normalized EPSC and IPSC traces for control (black) and deprived (magenta) neurons. Right: Mean E-I charge ratio for control (black) and deprived (magenta) neurons (control n = 12, deprived n = 12, P = 0.022, 2-sample t-test). **G**) Mean input resistance and capacitance for control (black) and deprived (magenta) neurons. **H**) Left: control intracortical (IC) versus thalamocortical (TC) E-I ratios (P = 0.84, 2-way t-test). Right: deprived intracortical versus thalamocortical E-I ratios (P = 4.1 x 10^-5^, 2-way t-test).

### Decreased intracortical E-I ratio can account for the drop in firing induced by brief MD

Given that thalamocortical and intracortical E-I ratios change in opposite directions after brief MD, we wanted to know if the net effect of these changes was to increase or decrease local layer 4 excitability. To model the changes we observed, we built a model network of recurrently coupled excitatory and inhibitory neurons (Figure 8A). We numerically simulated 5,000 leaky integrate-and-fire neurons (80% excitatory and 20% inhibitory) with sparse and random connectivity implemented as conductance-based synapses (Brunel, 2000). The neurons received independent excitatory spike trains with Poisson statistics with synaptic weight *j_s_* to model thalamic drive to a local network in layer 4 of V1. The strength of the thalamic drive to inhibitory neurons was stronger than the drive to excitatory pyramidal neurons (Figure 4), which we included through a multiplication of the overall input weight *j_s_* with a factor *g_fw_* > 1 for the drive to inhibitory neurons. To model input from other sources (such as intracortical feedback from neurons outside the local model circuit, or from higher cortical areas) neurons also received thalamocortical-independent excitatory spike trains. Input rates were chosen so that at baseline the networks generated excitatory and inhibitory firing rates as measured experimentally *in vivo* (Figure 8A, bar plot, Hengen et al., 2013). Thalamocortical firing rates were fixed across conditions, as mean firing rates of thalamic relay neurons in the LGN are reportedly not affected by acute lid suture (Linden et al., 2009).

**Figure 8:**
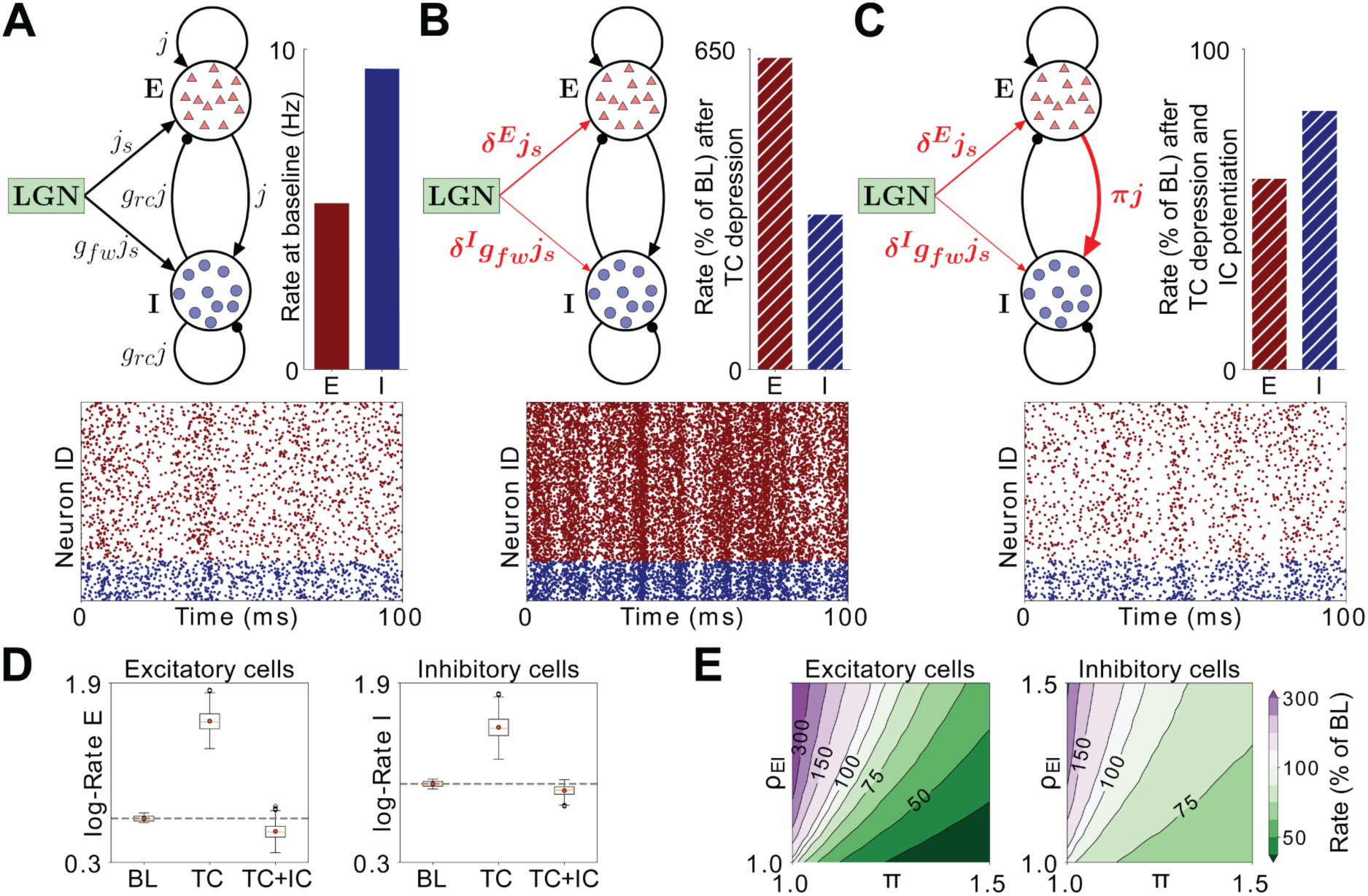
Firing rates in a model layer 4 circuit incorporating MD-induced synaptic changes. **A**) Schematic of the network model under baseline conditions. All excitatory synaptic weights in the network are given by the parameter *j*, and inhibitory connections are further scaled with a factor *g_rc_*. Excitatory neurons receive thalamic input with synaptic weight *j_s_*, while the stronger thalamic input onto inhibitory neurons is captured by multiplying *j_s_* with the factor *g_fw_* > 1. Bar plot: Firing rates of excitatory (E, red) and inhibitory (I, blue) neurons in the model at baseline (BL). Raster plot: Representative spiking activity across neurons over 100 ms at BL. **B**) Schematic of the network model after implementing synaptic depression in thalamocortical (TC) projections onto excitatory neurons (δ^E^) and greater depression onto inhibitory neurons (δ^I^), mirroring changes from Figures 3 and 4. Bar plot: firing rates as a percentage of BL resulting from these synaptic changes. Raster plot: representative spiking activity across neurons over 100 ms after implementing TC depression. **C**) Same as B), but with the addition of intracortical (IC) potentiation (π) of recurrent excitatory drive to inhibitory neurons (TC+IC). **D**) Box plots of the firing rates in 2,491 implementations of the network with randomly chosen parameters, simulated at BL, after TC depression and after TC depression and IC potentiation. The boxplots show the interquartile range (IQR) of the simulated firing rates (red symbol denotes the mean, red line denotes the median), with the whiskers denoting ± 1.5 IQR. Outliers (black circles) are firing rates beyond the ± 1.5 IQR. Rates are shown on a logarithmic scale for excitatory neurons (left panel) and inhibitory neurons (right panel). **E**) Combined effects of increased TC E-I ratio (ρ_EI_) and potentiation of recurrent excitatory drive to inhibition (π) on firing rates of excitatory (left panel) and inhibitory neurons (right panel). Firing rates are shown as percentage of BL for all combinations of π and ρ_EI_. Green-shifted color regions represent lower rates and purple-shifted regions represent higher rates relative to BL.

To test the impact of the E-I changes we observed following brief MD, we first modeled the observed change in thalamocortical E-I ratio (Figure 3C) by implementing a depression of thalamocortical strength onto excitatory neurons (δ^E^) and a larger depression of thalamocortical strength onto inhibitory neurons (δ^I^ < δ^E^). As expected, this resulted in increased firing of both cell types due to a net reduction in inhibitory drive (Figure 8B). Next, we added the observed shift in intracortical E-I ratio (Figure 7F), which was implemented through a potentiation of intracortical excitatory strength onto inhibitory neurons (π) (Maffei et al., 2006). We found that this additional change exerted a powerful effect that opposed the increase in thalamocortical E-I ratio, acting to decrease both excitatory and inhibitory firing rates, similar to the change seen *in vivo* following brief MD (Figure 8C, Hengen et al., 2013, 2016).

To ensure that these results were not sensitive to the specific choice of weights for overall recurrent connection strength (*j*) or the relative scale of recurrent inhibitory strength (*g_rc_*), we simulated 2,491 networks where we varied these parameters in a large range (Figure 8D, see also methods). Applying the synaptic changes as before generated consistent changes in firing rates across all networks, demonstrating a general effect of firing rate suppression that was not specific to the particular network parameters that we chose. Finally, to investigate whether this was specific to the observed values of E-I ratio or a more general phenomenon, we simulated networks with combinations of increased thalamocortical E-I ratios (ρ_EI_ = δ^E^/ δ^I^) and decreased intracortical E-I ratios (π). The model shows that π has a disproportionately larger effect on network firing compared to ρ_EI_ at a wide range of values (Figure 8E). Hence, our model suggests that the key factor driving a reduction in V1 excitability after brief MD is the shift in intracortical E-I ratio to favor inhibition.

## DISCUSSION

It has long been thought that LTD at thalamocortical synapses is the major or sole cause of the rapid MD-induced loss of visual responsiveness in V1 (Bear, 2003; Crozier et al., 2007; Espinosa and Stryker, 2012; Khibnik et al., 2010; Malenka and Bear, 2004; Yoon et al., 2009). Surprisingly, here we show that – despite confirming the induction of LTD at thalamocortical synapses in layer 4 - brief MD *increases* the E-I ratio of feedforward thalamocortical drive onto layer 4 pyramidal neurons, making the thalamocortical circuit more, not less, excitable. This increase in thalamocortical E-I ratio is driven by disproportionate depression of thalamocortical synapses onto PV+ interneurons, thus reducing the recruitment of feedforward inhibition in layer 4. In contrast, intracortical feedback E-I ratio onto layer 4 pyramidal neurons is reduced by brief MD, and a network model of layer 4 demonstrates that the net effect of these opposing changes in feedforward and feedback drive is to suppress firing in layer 4. Thus, feedforward and feedback E-I ratio onto the same postsynaptic target can be independently regulated by visual experience, and loss of visual responsiveness in layer 4 after brief MD during the critical period is due to a reconfiguration of intracortical circuits that leads to the recruitment of excess feedback inhibition.

### LTD and loss of visual responsiveness in layer 4

There is abundant evidence that brief MD during the critical period induces LTD at thalamocortical synapses onto layer 4 pyramidal neurons in both V1m and V1b (Crozier et al., 2007; Wang et al., 2013; Heynen et al., 2003). Manipulations that block this LTD also prevent the reduction in layer 4 visual evoked potentials (VEPs) elicited by stimulation of the deprived eye (Yoon et al., 2009), an observation that led to the conclusion that LTD of thalamocortical inputs onto layer 4 pyramidal neurons is the primary mechanism for loss of visual responsiveness in layer 4 (Smith et al., 2009). However, the measure of visual responsiveness (the rapid negative-going VEP) used in Yoon et al., 2009 primarily measures direct thalamocortical-evoked synaptic responses (Rahmat, 2009, Khibnik, et al., 2010), so is not expected to capture changes in intracortical circuitry or E-I ratio that might occur independently of thalamocortical LTD.

While we confirmed that brief MD induces LTD at thalamocortical synapses onto layer 4 pyramidal neurons, we used paired recordings to show that synapses onto PV+ interneurons are more strongly depressed than synapses onto nearby pyramidal neurons, suggesting that the reduced VEP following brief MD reflects depression at both synapse types. This is reminiscent of sensory deprivation-induced plasticity in barrel cortex (S1), where whisker plucking disproportionately reduces excitatory drive from layer 4 to layer 2/3 PV+ interneurons compared to layer 2/3 pyramidal neurons (House et al., 2011; Li et al., 2014). However, in S1 there is no net change in feedforward E-I ratio onto pyramidal neurons, due to concurrent potentiation of PV+ interneuron connections onto pyramidal neurons (House et al., 2011). Here, we find that brief MD in V1 induces disproportionate depression of thalamocortical strength onto PV+ interneurons with no concurrent potentiation of inhibitory strength. This in turn leads to an enhancement of thalamocortical-evoked E-I ratio onto layer 4 pyramidal neurons (and thus net disinhibition of the feedforward thalamocortical circuit) that our modeling suggests would produce an *increase* in network excitability if unopposed by other microcircuit changes. Thus, while our data are consistent with previous reports of MD-induced thalamocortical LTD and depression of VEPs (Crozier et al., 2007; Yoon et al., 2009), they cast doubt on the interpretation that LTD at thalamocortical synapses underlies the loss of visual responsiveness in V1 measured using extracellular spiking or intrinsic signal imaging (Kaneko et al., 2009, Hengen et al., 2013), both of which are sensitive to changes in intracortical circuitry. Instead, our data strongly support an alternative explanation in which intracortical inhibition is enhanced (relative to excitation) by brief MD, and this enhancement suppresses the excitability of the layer 4 microcircuit.

### Independent regulation of E-I ratio at feedforward and feedback circuits within layer 4

Many studies have examined the impact of sensory deprivation on PV+ fast-spiking (FS) inhibition within primary sensory cortex, using a variety of protocols to elicit and measure inhibition (Hengen et al., 2013; House et al., 2011; Kannan et al., 2016; Kuhlman et al., 2013; Maffei et al., 2006; Nahmani and Turrigiano, 2014). Together, these studies suggest that changes in intracortical inhibition are highly dynamic and depend critically on the length of sensory deprivation, the source of inhibition, and the neocortical layer under investigation (Hengen et al., 2013; Kannan et al., 2016; Kuhlman et al., Maffei et al., 2004, 2008, 2010). However, most studies have not separated feedforward from feedback inhibition, so the degree to which they can be independently regulated by sensory experience is unclear. In part, this is due to the difficulty in isolating these two intimately related aspects of inhibition; the same PV+ FS interneurons generate both feedforward and feedback inhibition within layer 4 and layer 2/3 (Figure 3A), and many methods of evoking inhibition (for example electrical or visual stimulation) will elicit both together. Here we have taken advantage of recent advances in mouse genetics and optogenetics to isolate and independently measure these two aspects of inhibition within layer 4. Unexpectedly, our data suggest that these two distinct inhibitory microcircuit motifs – one mediated by thalamocortical drive and the other mediated by recurrent pyramidal drive onto the same postsynaptic PV+ interneurons – change in opposite directions following brief MD.

Although PV+ FS synapses onto layer 4 pyramidal neurons are known to be plastic (Lefort et al., 2013; Maffei et al., 2004, 2006; Nahmani et al., 2014), changes at this synapse cannot account for either MD-induced increased feedforward or decreased feedback inhibition onto layer 4 pyramidal neurons, as we observed no change in either PV+ interneuron-to-pyramidal synaptic strength or in intrinsic excitability of PV+ interneurons. We probed putative unitary PV+ to pyramidal connections using ChR2 to activate individual labeled PV+ interneurons, a relatively high-throughput approach that allowed us to sample a large number of connections, and found that the distribution of synaptic weights was unaffected by brief MD. This result contrasts with a previous study that used paired recordings to demonstrate enhanced unitary connection strength at this synapse after MD from postnatal days 18-21 (Maffei et al., 2006). This discrepancy may reflect differences in the timing of MD, as postnatal day 18 is at the transition between the pre-critical period and the critical period proper (which opens around postnatal day 21); alternatively, it may reflect differences in sampling of the FS population, as Maffei et al. 2006 used firing properties to identify FS cells in rats, while here we used a mouse line with abundant labeling of PV+ FS cells in layer 4. Our data suggest that, rather than directly modulating inhibitory synaptic strength from PV+ interneurons to pyramidal neurons, brief MD modifies evoked inhibition by altering excitatory drive to inhibitory elements within layer 4.

Independent regulation of feedforward and feedback inhibition onto the same postsynaptic target requires either that the sources of inhibition differ, or that the same source of inhibition is differentially recruited by feedforward and feedback excitatory drive. While the relative reduction of feedforward inhibition we observed can be accounted for by strong depression of thalamocortical synapses onto PV+ interneurons, how intracortical feedback inhibition is enhanced relative to excitation is less clear. Both feedforward and feedback inhibition in layer 4 is largely mediated by PV+ interneurons. While SST+ interneurons (the other major GABAergic cell type in layer 4) do inhibit pyramidal neurons, they receive relatively weak thalamocortical drive (Beierlein et al., 2003; Cruikshank et al., 2010; Ji et al., 2016) and provide much stronger inhibition onto PV+ FS interneurons than onto layer 4 pyramidal neurons (Xu et al., 2013), and so they are thought to primarily *disinhibit* pyramidal neurons in layer 4. Plasticity of neocortical SST+ synapses is poorly explored and it is unknown whether sensory deprivation can modify the strength of these synapses onto either PV+ FS interneurons or pyramidal neurons (Scheyltjens and Archens, 2016; Urban-Ciecko and Barth, 2016), but reduced SST inhibition onto PV+ interneurons or increased SST inhibition onto pyramidal neurons are intriguing mechanisms that could explain enhanced intracortical inhibition onto pyramidal neurons following brief MD. Alternatively, potentiation of excitation from layer 4 pyramidal neurons to PV+ FS interneurons has been described following brief MD (Maffei et al., 2006) and provides another plausible mechanism for independently boosting intracortical inhibition without affecting feedforward thalamocortical recruitment of PV+ interneurons. For simplicity and consistency with known plasticity mechanisms, we modeled the change in intracortical E-I ratio as a change in excitatory drive from pyramidal to FS interneurons (Figure 8). In this model, the shift toward enhanced intracortical inhibition outweighs the effects of reduced thalamocortical inhibition over a wide range of network parameters; this is in large part because the vast majority of input to both PV+ interneurons and pyramidal neurons arises from recurrent excitatory synapses, so that even small changes to net intracortical excitatory drive to inhibitory neurons can have a powerful impact on layer 4 firing.

### Mechanisms of E-I regulation

Eyelid closure during MD leads to a degradation of retinal input to dLGN and the decorrelation of firing of dLGN relay neurons (Linden et al., 2009), and this decorrelation likely drives LTD by reducing the ability of thalamocortical afferents to effectively cooperate to depolarize postsynaptic pyramidal neurons (Smith et al., 2009). LTD has not previously been reported at thalamocortical synapses onto PV+ interneurons. LTD has been described at a number of other hippocampal and neocortical excitatory synapses onto PV+ interneurons (Lu et al., 2007; McMahan and Kauer, 1997; Nissen et al., 2010), but the pre-post activity patterns and signaling pathways required for its induction are variable and depend on circuit and cell type (Gibson et al., 2008; Lu et al., 2007). Our data suggest that the same MD-induced decorrelation of thalamocortical input that drives LTD at thalamocortical synapses onto pyramidal neurons is even more effective at driving LTD at synapses onto PV+ interneurons. This imbalance in LTD induction results in an enhanced feedforward excitability, which may serve to compensate for degraded sensory input by boosting the sensitivity of the system to the input that remains (Hennequin et al., 2017). An interesting question is whether this reconfiguration of the feedforward network is solely an early adaptive mechanism that eventually reverses, or whether it persists even in the face of prolonged sensory deprivation.

The cellular plasticity mechanisms that enhance feedback inhibition (relative to excitation) within the layer 4 recurrent circuit are less clear. Interestingly, unlike the change in feedforward inhibition (Figure 3C), the change in feedback inhibitory charge is primarily driven by a prolongation of inhibitory current rather than a change in peak current (Figure 7F). This suggests that the recruitment of inhibitory neurons by intracortical drive is prolonged after brief MD, presumably due to increased net excitation onto PV+ interneurons. An intriguing question is whether this plasticity in the feedback intracortical circuit is driven by preceding changes in feedforward drive. For instance, our modeling predicts that feedforward changes alone will enhance firing of both pyramidal neurons and PV+ interneurons and enhance synchrony (Figure 8B), which may be sufficient to drive plasticity at intracortical synapses onto inhibitory neurons (Caporele and Dan, 2008; Hennequin et al., 2017).

What purpose might it serve to boost the gain of incoming sensory drive, only to throttle it down again by suppressing recurrent excitability? One possibility is that these combined changes serve to enhance the signal-to-noise ratio within layer 4. The increased feedforward E-I ratio should boost the ability of thalamic inputs to drive pyramidal neurons, thus offsetting to some extent the impact of degraded retinal drive. Concurrently, the prolongation of recurrent inhibition is expected to prevent these signals from being amplified and spreading out in time and space. Thus, these changes together may constrain overall activity in layer 4 to prevent the amplification of noise, while still allowing the temporal pattern of thalamocortical input to influence layer 4 pyramidal spike patterns.

## ACKNOWLEDGEMENTS

Funding sources: NSF GRFP and NRSA F31 NS089170 (N.J.M.); R37NS092635 and R01EY025613 (G.G.T.); Max Planck Society (J.G. and L.M.A.R.).

## AUTHOR CONTRIBUTIONS

Conceptualization, N.J.M., G.G.T.; Methodology, N.J.M., L.M.A.R., J.G., G.G.T.; Software, N.J.M., L.M.A.R.; Formal Analysis, N.J.M., B.A.C., L.M.A.R.; Investigation, N.J.M., B.A.C.; Writing, N.J.M., L.M.A.R., J.G., G.G.T.; Visualization, N.J.M., L.M.A.R.; Supervision, J.G., G.G.T.; Funding Acquisition, N.J.M., J.G., G.G.T.

## DECLARATION OF INTERESTS

The authors declare no competing interests

## STAR METHODS

### CONTACT FOR REAGENT AND RESOURCE SHARING

Further information and requests for resources and reagents should be directed to the lead contact, Gina G. Turrigiano.

### EXPERIMENTAL MODEL AND SUBJECT DETAILS

#### Overview

Experiments were performed on Long-Evans rats (Figures 1, S1, 2, and S3), WT C57BL/6J mice RRID: IMSR_JAX:000664, (Figure 3), or various genetic mouse lines in C57BL/6J backgrounds (Figures 4-7 and S4, see below). For all experiments, both male and female pups between postnatal day (P) 25 and 28 were used for slice physiology. Litters were housed on a 12:12 light cycle with the dam (rats) or with a mating pair (mice) with free access to food and water. Pups were weaned and moved to a new cage with 2-4 littermates at P21. Wood chip-based cage bedding was replaced twice weekly, and animals were provided enrichment, including additional bedding material, plastic huts, and wooden gnawing blocks. All animals were housed, cared for, surgerized, and sacrificed in accordance with Brandeis IBC and IACAUC protocols.

#### Long-Evans Rats, Charles River Labs strain code: 006

Timed pregnant female Long-Evans rats were obtained from Charles River Labs (Wilmington, MA).

#### PV-Cre Mice, RRID: IMSR_JAX:008069

Mice carrying the PV-Cre allele express Cre recombinase in PV+ interneurons, without disrupting endogenous PV expression (Hippenmeyer et al., 2005). Mice homozygous for PV-Cre were crossed with mice homozygous for Rosa26-STOP-tdTomato (see below) to produce pups hemizygous for both alleles. These pups express tdTomato in PV+ interneurons (Figures 4, S4, and 5). Mice homozygous for PV-Cre were crossed with mice homozygous for Rosa26-STOP-ChR2-EYFP (see below) to produce pups hemizygous for both alleles. These pups express ChR2-YFP in PV+ interneurons (Figure 6).

#### Rosa26-STOP-tdTomato (Ai9) Mice, RRID: IMSR_JAX:007909

Mice carrying the Ai9 allele express tdTomato in a Cre recombinase-dependent manner (Madisen et al., 2010). Mice homozygous for Ai9 were crossed with mice homozygous for PV-Cre to produce pups hemizygous for both alleles. These pups express tdTomato in PV+ interneurons (Figures 4, S4 and 5).

#### Rosa26-STOP-ChR2-EYFP (Ai32) Mice, RRID: IMSR_JAX:012569

Mice carrying the Ai32 allele express ChR2-YFP fusion protein in a Cre recombinase-dependent manner (Madisen et al., 2012). Mice homozygous for Ai32 were crossed with mice homozygous for PV-Cre to produce pups hemizygous for both alleles. These pups express ChR2-YFP in PV+ interneurons (Figure 6).

#### Scnn1a-Tg3-Cre Mice, RRID: IMSR_JAX:009613

Mice carrying the Scnn1a-Tg3-Cre allele express Cre recombinase in primary sensory cortical layer 4 pyramidal neurons (Madisen et al., 2010). Mice hemizygous for Scnn1a-Tg3-Cre were crossed with WT C57BL/6J mice to produce 50% of pups hemizygous for Scnn1a-Tg3-Cre. Pups hemizygous for Scnn1a-Tg3-Cre were intracortically injected with AAV-FLEX-ChR2-tdTomato so that a sparse population of layer 4 pyramidal neurons within V1m expressed ChR2-tdTomato (Figure 7).

### METHOD DETAILS

#### Anesthesia

For virus injection and lid suture surgeries (below), animals were briefly anesthetized in a sealed container containing 3-4 % isoflurane in air, followed by subcutaneous injection with a cocktail containing ketamine, xylazine, and acepromazine (mass ratio 70 : 5 : 1 for mice, 70 : 3.5 : 0.7 for rats). Depth of anesthesia was continuously monitored via toe pinch response and respiratory rate during surgery, and drug cocktail was re-administered in increments of 25% of original dose as necessary to maintain surgical anesthesia.

#### Virus injections into dLGN and V1m

P12-P16 animals were anesthetized as above. Fur covering the scalp was shaved with electric hair clippers, and the scalp was cleaned with isopropyl alcohol followed by application of betadine (iodine solution). Eyes were covered with sterile ophthalmic ointment to prevent desiccation during surgery. Animals were mounted on a stereotaxic frame, and the skull was exposed with a single incision such that bregma and lambda were visible. Dorsal LGN (Figures 1-4, S1, S3, and S4) or V1m (Figure 7) were targeted using stereotaxic coordinates obtained from brain atlases after adjusting for the lambda-bregma distance for age. Small regions of the skull were thinned bilaterally using a dental drill and then removed carefully with fine forceps under a dissection microscope. Virus was diluted 1/5 - 1/20 original titer with sterile saline prior to injection. A glass micropipette pulled to a fine point delivered 500 nL (rat dLGN), 200 nL (mouse dLGN), or 75 nL (mouse V1m) of virus-containing solution at the targeted depth. The skull was flushed with sterile saline and the scalp was sutured shut with 6-0 nylon sutures. Animals were rehydrated following surgery with a single subcutaneous injection of warm sterile saline (5% body weight), and simultaneously a single dose of penicillin was delivered to prevent infection. Animals recovered on a heating pad for 8-16 hr with free access to water and soft food and were returned to their litters afterwards. Meloxicam was delivered subcutaneously following the surgery and for 2-3 days after to reduce pain and inflammation. Antibiotic ointment and 2% lidocaine jelly were also applied to the sutured region for 2-3 days following surgery to prevent infection, desiccation, and pain at the sutured region.

#### Lid sutures

P22-P23 pups were anesthetized as above. One eye was chosen randomly for lid suture, and the other was covered in sterile ophthalmic ointment to prevent desiccation during surgery. The eye to be surgerized was first cleaned with betadine, followed by flushing with sterile saline. 1-2 mm of the lower part of each eyelid was carefully trimmed using fine surgical scissors, followed by a second flushing of the eye with sterile saline. The eyelids were then sutured closed with 3 6-0 nylon or silk sutures. Lastly, the sutured eyelid was covered in antibiotic ointment to prevent infection. Meloxicam was delivered subcutaneously following the surgery and for 2 days after to reduce pain and inflammation. Sutures were checked each day and animals were not used if sutures were not intact.

#### Ex vivo acute brain slice preparation

Coronal brain slices (300 µm) containing V1 were obtained from the control and deprived hemispheres of each animal, using a procedure modified from previous studies (Lambo and Turrigiano, 2013; Lefort et al., 2009; Maffei et al., 2006) and included ‘protective recovery’ immediately post-slicing (Ting et al., 2014). Animals were anesthetized with isoflurane deeply enough to have no response to firm toe pinch, followed by decapitation and whole brain removal. A coronal section normal to the midline was carefully blocked in a single plane from frontal cortex using a scalpal, and the brain was mounted to a slicing chamber via cyanoacrylate adhesive attached to this exposed surface. Slices were immediately cut on a Leica VT1000S vibratome in chilled (1 °C) carbogenated (95 % O_2_, 5 % CO_2_) standard ACSF containing (in mM): 126 NaCl, 25 NaHCO3, 3 KCl, 2 CaCl2, 2 MgSO4, 1 NaH2PO4, 0.5 Na-Ascorbate, with dextrose added to bring osmolarity to 315 mOsm, and titrated with HCl to bring pH to 7.35. Slices were then immediately transferred to a warm (34 °C) chamber filled with a continuously carbogenated ‘protective recovery’ choline-based solution containing (in mM): 110 Choline-Cl, 25 NaHCO3, 11.6 Na-Ascorbate, 7 MgCl2, 3.1 Na-Pyruvate, 2.5 KCl, 1.25 NaH2PO4, and 0.5 CaCl2, with dextrose added to bring osmolarity to 315 mOsm, and titrated with HCl to bring pH to 7.35. Following 10 min of incubation, slices were then transferred to warm (34 °C) carbogenated standard ACSF and incubated another 20 min, before being taken out of the incubator and allowed to return to room temperature. Slices were used for electrophysiology between 1-6 hr post-slicing.

#### Electrophysiology

##### Identification of V1m, layer 4, and cell types

V1m was identified in acute slices using the rat brain atlas (Paxinos and Watson 1998) and mouse brain atlas (Paxinos and Franklin 2001), using the shape and morphology of the white matter and hippocampal formation as a reference. V1 was also identified using several other criteria, such as a conspicuous high contrast band corresponding to layer 4, as well as thalamocortical afferent fluorescence (Figures 1-4, S1, S3, and S4), high density of PV+ interneurons (Figures 4-6, and S4), and Scnn1a+ pyramidal neurons (Figure 7), with V1m corresponding to the medial most section of this region. Pyramidal neurons were identified by the presence of an apical dendrite and teardrop shaped soma, and morphology was confirmed by post hoc reconstruction of biocytin fills, with apical dendrites reaching layer 1. PV+ interneurons were identified by reporter expression driven by the PV-Cre allele (see above).

##### Pipettes and internal solutions

Borosilicate glass recording pipettes were pulled using a Sutter P-97 micropipette puller, with acceptable tip resistances ranging from 2-5 MΩ. For Figures 1, 2, and 5, K+ Gluconate-based internal recording solution was modified from Lambo and Turrigiano, 2013, and contained (in mM) 100 K-gluconate, 10 KCl, 10 HEPES, 5.37 Biocytin, 10 Na-Phosphocreatine, 4 Mg-ATP, and 0.3 Na-GTP, with sucrose added to bring osmolarity to 295 mOsm and KOH added to bring pH to 7.35. For Figures 3-4 and 6-7, Cs+ Methanesulfonate-based internal recording solution was modified from Xue et al., 2014, and contained (in mM) 115 Cs-Methanesulfonate, 10 HEPES, 10 BAPTA•4Cs, 5.37 Biocytin, 2 QX-314 Cl, 1.5 MgCl_2_, 1 EGTA, 10 Na_2_-Phosphocreatine, 4 ATP-Mg, and 0.3 GTP-Na, with sucrose added to bring osmolarity to 295 mOsm, and CsOH added to bring pH to 7.35.

##### Acquisition hardware, software, and optics

All recordings were performed on submerged slices, continuously perfused with carbogenated 34 °C recording solution. Neurons were visualized on an Olympus BX51WI upright epifluorescence microscope using a 10x air (0.13 numerical aperture) and 40x water-immersion objective (0.8 numerical aperture) with infrared-differential interference contrast optics and an infrared CCD camera. Up to 3 neurons were simultaneously patched using pipettes filled with internal as described above, well-chlorided silver electrodes, and MPC-200 micromanipulators (Sutter Instrument, Novato CA). Data were low-pass filtered at 6 kHz and acquired at 10 kHz (except for data in Figures 1 and S1, which were low-pass filtered at 2.6 kHz and acquired at 5 kHz) with Multiclamp 700B amplifiers and CV-7B headstages (Molecular Devices, Sunnyvale CA) and an analog-to digital data acquisition system (National Instruments, Woburn MA). Data was acquired using an in-house program written in Igor Pro (Wavemetrics, Lake Oswego OR), and all post-hoc data analysis was performed using in-house scripts written in MATLAB (Mathworks, Natick MA). For optogenetic photostimulation, 473 nm blue light was emitted from a 50 mW DPSS laser, which was gated by a LS2 2 mm uni-stable shutter (Vincent Associates, Rochester NY), and light intensity was controlled with a NDM2 filter wheel (Thor Labs, Newton NJ). Laser light was routed through the optical path of the microscope and focused to a ∼50 µm diameter spot through the 40x objective.

##### Thalamocortical quantal amplitudes (Figure 1)

For all data here and below, control and deprived were obtained from the same animals, with deprived data obtained from V1m contralateral to the deprived eye, and control data obtained from ipsilateral V1m. For mEPSC and evoked quantal thalamocortical recordings, layer 4 pyramidal neurons were voltage clamped to −70 mV in standard ACSF containing a drug cocktail of TTX (0.2 µM), APV (50 µM), picrotoxin (25 µM), and 4-AP (100 µM) with a K+ Gluconate-based internal solution. 20 s traces were obtained under high gain (10-50x). Baseline traces were first obtained (for spontaneous mEPSCs) and then traces were obtained while stimulating continuously with 473 nm laser light for 2-5 s. Stimulus intensity was chosen such that evoked frequency was well above spontaneous frequency, but individual events were still clearly visible. Event inclusion criteria included amplitudes greater than 5 pA and rise times less than 3 ms. Neurons were excluded from analysis if series resistance was above 20 Mohms or if resting membrane potential was above −50 mV. Quantal data were analyzed in a fully automated manner (except for frequency data in Figure S1A, which involved manual analysis) using in-house scripts written in MATLAB (Mathworks, Natick MA).

##### Thalamocortical LTD induction (low frequency stimulation, Figure 2)

A low frequency stimulation (LFS) LTD induction paradigm was adapted from Crozier et al., 2007 for use with optogenetic stimulation of thalamocortical afferents. Whole cell voltage clamp recordings were obtained from up to 3 layer 4 pyramidal neurons simultaneously in V1m in standard ACSF with a K+ Gluconate-based internal solution, and cells were held at −70 mV. Laser stimulus strength was adjusted until 2 ms pulses delivered once per minute evoked thalamocortical EPSCs between 100-800 pA. Thalamocortical EPSCs were evoked once per minute for a 10 minute baseline period. LFS LTD was induced in voltage clamp by pairing 1 Hz optogenetic stimulation with a brief (100 ms) postsynaptic-step depolarization from −70 mV to −50 mV for each of 600 total pulses. Following LFS LTD induction, Thalamocortical EPSCs were again evoked once per minute for a 45 min post-induction period. For quantification in Figure 2E-G, post-induction amplitudes represent response for each cell average over the 5-30 min following end of LTD induction. Data were discarded if resting membrane potential was above −55 mV or if series resistance changed more than 25% during the recording.

##### E-I ratios (Figures 3, S3, and 7)

Experimental procedure for measurement of E-I ratio was adapted from Xue et al., 2014. Whole cell voltage clamp recordings were obtained from up to 3 layer 4 pyramidal neurons simultaneously in V1m in standard ACSF with a Cs+ Methanesulfonate-based internal solution. EPSCs and IPSCs were obtained optogenetically by stimulating either thalamocortical afferents (Figure 3 and S3) or other nearby pyramidal neurons (Figure 7) once per minute while alternating between the reversal potential of excitation and inhibition for each stimulus. In order to experimentally determine the reversal potential for inhibition, we used mice with both PV-Cre and Rosa26-STOP-ChR2-EYFP alleles, which express ChR2-YFP in PV+ interneurons. We recorded from layer 4 pyramidal neurons while stimulating groups of nearby ChR2-YFP+ PV+ interneurons. The holding potential was stepped up or down after each stimulus until the resulting IPSC amplitude was as close as possible to 0. In order to experimentally determine the reversal potential for excitation, we perfused a drug cocktail of TTX (0.2 µM), APV (50 µM), picrotoxin (25 µM), and 4-AP (100 µM) in standard ACSF while recording from layer 4 pyramidal neurons and optogenetically stimulating either thalamocortical afferents or nearby ChR2+ pyramidal neurons. The holding potential was stepped up or down after each stimulus until the resulting EPSC amplitude was as close as possible to 0. The measured inhibitory and excitatory reversal potentials were close to −70 mV and 0 mV after adjusting for the liquid junction potential. For thalamocortical E-I ratio, minimum acceptable EPSC and IPSC amplitudes were 150 pA and 600 pA, and for intracortical E-I ratio, minimum acceptable EPSC and IPSC amplitudes were 50 pA and 600 pA. Any traces that contained obvious polysynaptic activity were discarded. Recordings were excluded if series resistance was greater than 25 MΩ.

##### Paired pyramidal / PV+ interneuron thalamocortical EPSCs (Figure 4)

A PV+ interneuron and nearby layer 4 pyramidal neuron were patched and voltage clamped to −70 mV in standard ACSF containing a drug cocktail of TTX (0.2 µM), APV (50 µM), picrotoxin (25 µM), and 4-AP (100 µM), which guaranteed that thalamocortical EPSCs were monosynaptic and not contaminated with IPSCs. Cs+ Methanesulfonate-based internal was used for this experiment. Stimulus laser intensity was increased until 2 ms pulses evoked EPSC amplitudes greater than 100 pA in both cells. Thalamocortical EPSCs were evoked once per minute at a range of stimulus intensities. Recordings were excluded if series resistance was greater than 30 MΩ, or if there was greater than 30% difference in series resistance values between pyramidal neuron and PV+ interneuron within a pair.

##### Paired pyramidal / PV+ interneuron thalamocortical-evoked firing (Figure S4)

A PV+ interneuron and nearby layer 4 pyramidal neuron were patched and held in current clamp in standard ACSF, and a small dc bias current was injected to maintain resting membrane potential at −70 mV. K+ Gluconate-based internal was used for this experiment. A ramp of stimulus laser intensities was delivered, with 2 ms pulses delivered each minute. Recordings were excluded if resting membrane potential was greater than −50 mV.

##### Intrinsic excitability measurement (Figure 5)

Whole cell recordings were performed on layer 4 PV+ interneurons or pyramidal neurons in current clamp mode in standard ACSF containing a drug cocktail of APV (50 µM), DNQX (25 µM), and picrotoxin (25 µM), with K+ Gluconate-based internal solution. A small dc bias current was injected to maintain resting membrane potential at −70 mV. Then, 1 s long current injections of increasing magnitudes were delivered to generate F-I curves. For PV+ interneurons, current injections were increased in increments of 20 pA until 500 pA, after which they were increased in increments of 100 pA. For pyramidal neurons, current injections were increased in increments of 20 pA. Recordings were excluded if series resistance was greater than 25 MΩ. For pyramidal neurons, recordings were excluded if input resistance was less than 100 MΩ or if resting membrane potential was greater than −60 mV. For PV+ interneurons, recordings were excluded if input resistance was less than 40 MΩ or if resting membrane potential was greater than −50 mV.

##### Optogenetically evoked unitary IPSCs (PV+ interneuron minimal stimulation, Figure 6)

Whole cell recordings were simultaneously performed on up to 3 neighboring layer 4 pyramidal neurons voltage clamped at 0 mV in standard ACSF with Cs+ Methanesulfonate-based internal solution. Using epifluorescence, individual nearby ChR2-YFP+ PV+ interneurons were targeted with low enough 473 nm laser stimulation intensity such that no evoked IPSCs were visible with 2 ms pulses delivered every 30 s. Then, laser stimulus intensity was slowly increased until a stimulus intensity was reached where roughly 50% of stimuli resulted in an evoked IPSC in one or more of the recorded pyramidal neurons. Other indicators of evoked unitary IPSCs included consistent amplitude and waveform, as well as a tendency for all of the recorded pyramidal neurons to simultaneously exhibit response or failure. Recordings were excluded if series resistance was greater than 25 MΩ.

##### Modeling layer 4 network and changes due to E-I ratio

The simulated model network consisted of 5,000 leaky integrate-and-fire neurons, 80% of which are excitatory and 20% inhibitory (Brunel, 2000). The membrane potential of each neuron was described by:

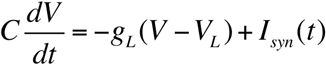

where *I_syn_ (t)* denotes the synaptic input into a neuron

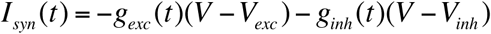

and *g_exc_ (t)* the total excitatory and *g_inh_ (t)* the total inhibitory conductance. The excitatory and inhibitory reversal potentials were *V_exc_* and *V_inh_*, respectively. Each cell received a total of 400 excitatory and 100 inhibitory connections from randomly chosen cells within the local circuit (probability of two cells being connected is 10%). Connections between neurons were implemented as conductance-based synapses with an exponential kernel, where upon arrival of a spike, the conductance of the postsynaptic cell increased by a fixed amount (“peak conductance”) and afterwards decayed exponentially with a time constant of 5 ms. All excitatory connections within the circuit had the same peak conductance *j* and all inhibitory connections had the same peak conductance *g_rc_j*.

All neurons in the circuit received external excitatory drive from thalamus, modeled as 100 uncorrelated Poisson spike trains with rate *R_LGN_* = 5 Hz. The peak conductance for thalamic drive to excitatory cells was given by the parameter *j_s_*. The strong feedforward drive to inhibitory cells, as seen in electrophysiological experiments (Figure 4), was included in the model as a pre-factor *g_fw_* to the peak conductance of thalamic inputs to inhibition. In addition to this thalamic input, both excitatory and inhibitory cells received 100 Poisson spike trains with rate *R_BKG_* = 14.2 Hz from an unspecified source (“background”). This could denote input from microcircuits in layer 4 farther from the one being modeled, or input from other layers. The peak conductance of these inputs to both cell types was the same for the baseline (BL), thalamocortical (TC) and thalamocortical and intracortical (TC+IC) scenarios. All circuit and single neuron parameters are summarized in Table 1. This model circuit was simulated for 9 s to estimate the average population firing rate in the steady state in all three cases BL, TC and TC+IC (Figure 8A-C). All network simulations were performed using NEST (Gewaltig and Diesmann, 2007).

**Table 1.** Parameters of the neurons and network model (Figure 8A-C, E).

| Parameter | Variable | Value |
| --- | --- | --- |
| Total number of cells | $N$ | 5,000 |
| Number of excitatory cells | $N_E$ | 4,000 |
| Number of inhibitory cells | $N_I$ | 1,000 |
| Connection density/probability | $\varepsilon$ | 0.1 |
| Capacitance | $C$ | 200 pF |
| Excitatory peak conductance | $j$ | 0.3 nS |
| Inhibitory peak conductance | $g_{rc}j$ | 2.4 nS |
| Leak conductance | $g_L$ | 10 nS |
| Excitatory reversal potential | $V_{exc}$ | 0mV |
| **Inhibitory reversal potential** | $V_{inh}$ | **-85 mV** |
| **Leak reversal/resting potential** | $V_L$ | **-70 mV** |
| **Firing threshold** | $V_{\Theta}$ | **-50 mV** |
| **Thalamic/background input weight** | $j_s$ | **0.5 nS** |
| **Inhibitory thalamic input weight** | $g_{fw}j_s$ | **0.625 nS** |
| **Thalamic input rate** | $R_{LGN}$ | **8 Hz** |
| **Background input rate** | $R_{BKG}$ | **14.2 Hz** |
| **Number of thalamic input synapses** | $n_{LGN}$ | **100** |
| **Number of background input synapses** | $n_{BKG}$ | **100** |

To test the generality of the modeling results to the choice of parameters, we used a Monte-Carlo approach. We simulated 100,000 implementations of the network with randomly chosen parameters: excitatory coupling scale *j*, the relative scaling of recurrent inhibition *g_rc_*, and the rates of the two input sources, thalamus (*R_LGN_*) and background (*R_BKG_*). For each network implementation, all four parameters were chosen randomly within a certain interval (see Table 2). From all 100,000 implementations, we selected 2,491 networks where the firing rates in BL were matched to firing rates measured in vivo before MD (Hengen et al., 2013), allowing for ± 10% deviations around the measured firing rates. Each selected network underwent the same plastic changes corresponding to the TC and TC+IC scenarios, with the outcome summarized in Figure 8D.

**Table 2.** Ranges of varied parameters in the Monte-Carlo simulation (Figure 8D).

| **Parameter** | **Minimum** | **Maximum** |
| --- | --- | --- |
| Coupling scale $j$ | 0.1 nS | 0.4 nS |
| Relative inhibitory strength $g_{rc}$ | 7 | 10 |
| Thalamic input rate | 5 Hz | 15 Hz |
| Background input rate | 5 Hz | 15 Hz |

To further investigate the interaction of the measured opposite shifts in thalamocortical and intracortical E-I ratios, we simulated a single implementation of the network on a grid of combinations of the two E-I ratios. The depression of thalamic inputs to excitatory cells was fixed to the measured value at early MD (90% of control hemisphere). This amount of depression was combined with varying amounts of depression of thalamic inputs to inhibitory cells, ranging from 90% down to 60% of their baseline value. The depression of thalamocortical synapses onto inhibitory cells was then combined with the depression of drive to excitatory cells into a single parameter *ρ_EI_* = *δ_E_/δ_I_*, which accounts for the change in the thalamocortical E-I ratio. With the chosen values for *δ_I_*, *ρ_EI_* varied between 1 and 1.5 (y-axis in Figure 8E). All of the increased thalamocortical E-I ratios were then paired with decreased intracortical E-I ratios, which were implemented as potentiation of the drive from local excitatory cells onto inhibitory cells. This increase of excitatory drive to inhibition, accounting for the decrease in intracortical E-I ratio, was given by the parameter π (x-axis in Fig. 8E) and varied in the same range as *ρ_EI_*. The firing rates for each pair (*ρ_EI_, π*) were compared relative to the BL scenario *ρ_EI_* = *π* = 1 (Figure 8E).

### QUANTIFICATION AND STATISTICAL ANALYSIS

All data analysis was performed using in-house scripts written in MATLAB (Mathworks, Natick MA). For each experiment, means ± SEM were provided within the results section of the text, and n’s, P values, and statistical tests were provided within figure captions. For normally distributed data, a 2-sample t-test was used to assess P value. For data that were clearly non-normally distributed, a Wilcoxon rank sum test was used to assess P value. Lastly, for comparisons between distributions of data, a two-sample Kolmogorov-Smirnov test was used to assess P value. Significance was defined as P < 0.05.

### DATA AND SOFTWARE AVAILABILITY

Description: URL/supplemental file name goes here

