## Supplementary Materials for "Sensory Deprivation Independently Regulates Neocortical Feedforward and Feedback Excitation-Inhibition Ratio"

***Figure S1: Evoked thalamocortical quantal events and spontaneous mEPSCs show similar amplitude distributions and event kinetics.*** **A**) Spontaneous mEPSC frequency (grey) and evoked mEPSC frequency (black) for a random subset of manually analyzed neurons, with lines connecting spontaneous and evoked values for each neuron (n = 20 neurons, P = $5.5 \times 10^{-19}$, 2-sample t-test). **B**) Mean normalized event amplitude histogram for all neurons, with stimulus-evoked thalamocortical quantal events plotted in black and spontaneous mEPSCs in grey. **C**) Peak-scaled mean event waveforms for thalamocortical quantal events (black) and spontaneous mEPSCs (grey), to illustrate kinetics. **D**) Spontaneous mEPSC amplitudes measured for the same neurons as in Figure 1, with control in black and deprived in magenta (control n = 22, deprived n = 18, P = 0.26, 2-sample t-test).

***Figure S3: Brief MD increases thalamocortical-evoked E-I ratio in Long-Evans rats during the critical period.*** **A**) Mean EPSC charge-normalized E and I traces for control (black) and deprived (magenta) neurons. **B**) Mean E-I charge ratio for control (black) and deprived (magenta) neurons. (control n = 9, deprived n = 13, P = 0.032, 2-sample t-test).

***Figure S4: PV+ interneurons readily fire to thalamocortical stimulation, whereas pyramidal neurons do not.*** **A**) Example traces from a paired recording between a PV+ interneuron (blue) and nearby pyramidal neuron (red) showing firing responses to low, medium, and high intensity optogenetic stimulation of thalamocortical afferents. **B**) Mean evoked spikes for all PV+ interneurons (blue) and pyramidal neurons (red) versus laser stimulus strength (n = 6 pairs, P = $2.3 \times 10^{-42}$ two-sample Kolmogorov-Smirnov test).
