## Supplementary figures and images for "Sensory Deprivation Independently Regulates Neocortical Feedforward and Feedback Excitation-Inhibition Ratio"

### Supplementary Materials

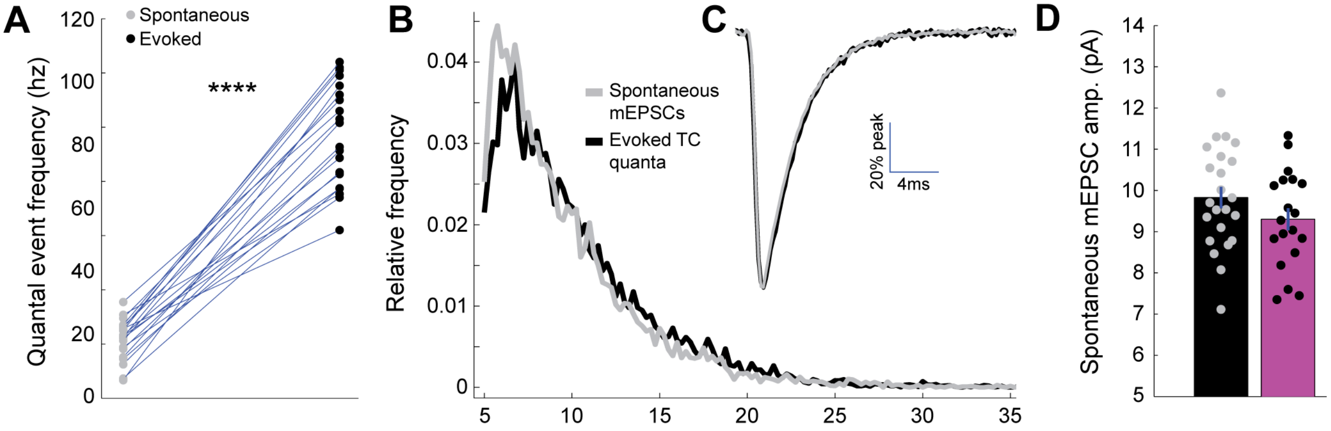

### Supplementary Materials

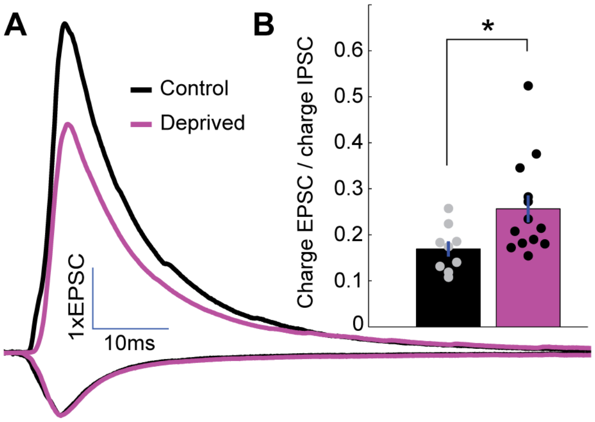

### Supplementary Materials

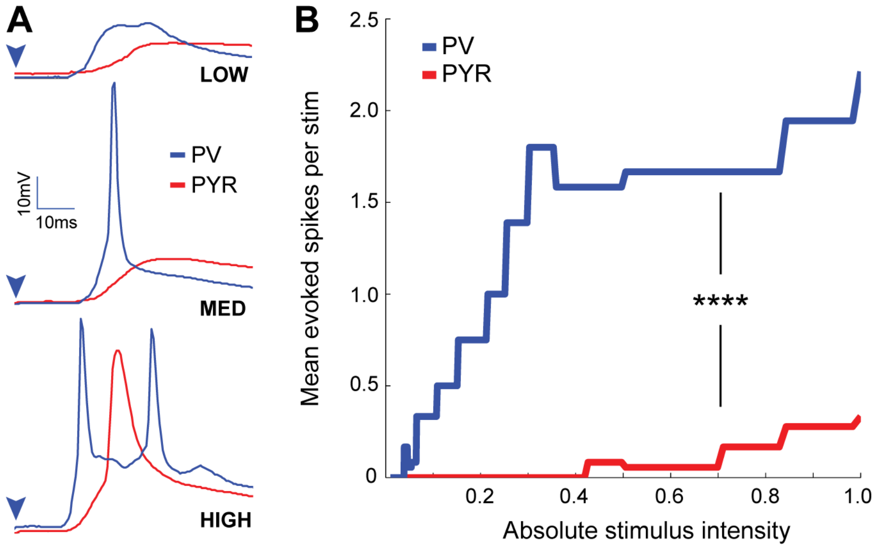
